# Hubness reduction improves clustering and trajectory inference in single-cell transcriptomic data

**DOI:** 10.1101/2021.03.18.435808

**Authors:** Elise Amblard, Jonathan Bac, Alexander Chervov, Vassili Soumelis, Andrei Zinovyev

**Author notes:** These authors contributed equally to this work (Elise Amblard, Jonathan Bac).

## Abstract

**Background:** Single-cell RNA-seq datasets are characterized by large ambient dimensionality, and their analyses can be affected by various manifestations of the dimensionality curse. One of these manifestations is the hubness phenomenon, i.e. existence of data points with surprisingly large incoming connectivity degree in the neighbourhood graph. Conventional approach to dampen the unwanted effects of high dimension consists in applying drastic dimensionality reduction. It remains unexplored if this step can be avoided thus retaining more information than contained in the low-dimensional projections, by correcting directly hubness.

**Results:** We investigate the phenomenon of hubness in scRNA-seq data in spaces of increasing dimensionality. We also link increased hubness to increased levels of dropout in sequencing data. We show that hub cells do not represent any visible technical or biological bias. The effect of various hubness reduction methods is investigated with respect to the visualization, clustering and trajectory inference tasks in scRNA-seq datasets. We show that hubness reduction generates neighbourhood graphs with properties more suitable for applying machine learning methods; and that it outperforms other state-of-the-art methods for improving neighbourhood graphs. As a consequence, clustering, trajectory inference and visualisation perform better, especially for datasets characterized by large intrinsic dimensionality.

**Conclusion:** Hubness is an important phenomenon in sequencing data. Reducing hubness can be beneficial for the analysis of scRNA-seq data with large intrinsic dimensionality in which case it can be an alternative to drastic dimensionality reduction.

## Introduction

The technology of single-cell omics profiling revolutionized many fields of modern molecular biology, providing more direct ways to study such biological phenomena as differentiation,^1^ development,^2^ heterogeneity of cancer cell populations and related resistance to treatment.^3, 4^ However, the analysis of single-cell datasets remains challenging, amplifying the difficulties already encountered in the analysis of bulk omics measurements as well as introducing new ones, specific to single-cell technologies.^5, 6^

From the geometrical point of view, a set of single-cell measurements can be represented as a cloud of data points, where each point represents a cell. This cloud is embedded in a space with overwhelmingly large dimensionality: thus, tackling only the case of scRNA-seq, a typical single-cell dataset provides information about measurable variability of up to twenty thousand gene expressions. Hence, the formal (or ambient) dimension of the data space is close to 10^4^ by order of magnitude. Due to the underlying biological mechanisms, the expression profiles of individual genes are coupled through complex network of linear and non-linear dependencies. This makes the effective or intrinsic dimensionality (ID) of the data point cloud much lower than the ambient one, even if the number of cells greatly exceeds 10^4^. For example, if all genes were perfectly linearly correlated to one common latent factor, then all cells would be located on a single line segment embedded in the multi-dimensional space, and the value of intrinsic dimensionality would be one, independently of the ambient dimensionality. Real gene expression datasets are influenced by more than one latent factor, and, intuitively, the number of distinct latent factors corresponds to the global intrinsic dimensionality of the data.^7^ The estimates of the intrinsic dimensionality of single-cell datasets can vary from 3-4 to hundreds.^8, 9, 4^

The difficulties of dealing with many dimensions in the data analysis are broadly referred to as the “curse of dimensionality”.^10^ Talking about the curse becomes relevant when the logarithm of the number of data points is less than the intrinsic dimensionality of the data.^11^ In practice, it means that certain manifestations of the dimensionality curse might appear starting with an intrinsic dimensionality as low as 10. These manifestations are various: among the most known is the concentration of distances quantified as the contrast between “close” and “far” distances in a dataset. Several approaches were proposed to compensate for the undesirable effect of distance concentration.^10, 12, 13^ However, in practice, it was demonstrated that it can not be avoided by global modifications of data space metrics.^14^ Another manifestation of the high-dimensional geometry is almost perfect linear separability of the data point cloud.^15^

In most of the presently used analysis workflows, single-cell datasets are subjected to drastic dimensionality reduction before applying many unsupervised machine learning methods.^16, 17^ After applying such dimensionality reduction, the ambient dimensionality is mechanistically truncated even if the intrinsic dimensionality of the data was higher (in this case, it is truncated as well). Indeed, this common practice aims at reducing possible manifestations of the curse of dimensionality such as distance concentration, at the cost of neglecting signals that are potentially contained in higher dimensions. Typical reduced dimensionality used to construct the k-NN graphs equals to 20-30. Moreover, for trajectory inference, the dimensionality of single-cell data is frequently reduced to 2 or 3.^8, 18^ This is in striking contrast with the observation that in a typical scRNA-seq dataset, first few tens of principal components explain only a small fraction of total variance (5-10%), unlike in case of bulk transcriptomic. The question of whether this is a mere consequence of the dropout in single-cell measurements or of an indeed intrinsically high-dimensionality of scRNA-seq data, has been poorly studied so far. Also, it remains unclear if the practice of reducing the dimension of single-cell data could eliminate useful information, and whether it is the only way to fight the dimensionality curse.

Yet another manifestation of the curse of dimensionality, much less known and discussed, is the hubness phenomenon. It has been described that in highdimensional settings some points might be surprisingly popular among the k-Nearest Neighbors (k-NN) of other points. This observation was formalized in,^19^coining the term “hub” to describe those points. In a more formal manner, hubness of a data point is the indegree of the corresponding node in the k-NN graph which reflects the number of times it appears among all the k-NN lists of other points. When looking at the distribution of hubness score as a function of the dimension of the data, Radovanovic et al. observed that part of the distribution can shift to the right when the dimension increases, forming a fat tail (Supplementary Figure 1A). The points in this tail have larger hubness scores compared to the rest of the distribution: they are designated as hubs. The tail of the distribution frequently follows a power law of Lévy, typical of many scale-free networks (Supplementary Figure 1B).

Dealing with the hubness phenomenon is crucial when computing and exploiting k-NN graphs, which is the most important ingredient of most of the currently used computational approaches for single-cell data analysis.^16, 20^ Simple or modified k-NN graphs are used for the tasks of clustering, non-linear dimensionality reduction, trajectory inference in single-cell datasets. Presence of hubs in the k-NN graph impacts their expected, from low-dimensional intuition, properties: for example, it can greatly impact the structure of geodesic distances along the graph between individual cells. Hubs (and antihubs, i.e. data points which are not nearest neighbours of any other point) make the structure of k-NN graphs heterogeneous in terms of connectivity which can violate the underlying assumptions for meaningful application of, for example, graph-based clustering algorithms. It is surprising that the hubness properties of the neighbourhood graphs produced from single-cell data and the impact of hubness on downstream analyses has never been studied so far.

Of note, in the popular non-linear dimensionality reduction approach Uniform Manifold Approximation and Projection (UMAP), weights of the neighbourhood graph edges, being transformations of the multidimensional distances, are introduced such that for each local neighbourhood, they would be characterized by sufficient contrast.^21^ As a consequence, the neighbourhood graph exploited by UMAP, is expected to have more regular properties (e.g. more uniform connectivity) than the standard k-NN graph, when computed in data spaces characterized by large intrinsic dimensionality. Due to this feature, UMAP method does not require the preliminary dimensionality reduction step, unlike many other methods, in its applications to single-cell data. However, it remains unclear if this way of constructing the k-NN graphs is the only one and how much it can be improved.

Hubness reduction methods aim at explicitly reducing the hubness of the k-NN graph, usually through specific transformations of the distance matrix.^22^ Interestingly, hubness reduction can be used instead of dimensionality reduction, in order to compensate for certain manifestations of the dimensionality curse. In this study we hypothesized that application of hubness reduction methods can be beneficial whenever performing single-cell data analyses using k-NN or other neighbourhood graphs or distance matrices as an essential ingredient. We explicitly and systematically evaluate the effect of hubness reduction on clustering, trajectory inference and visualisation in single-cell datasets, using previously established benchmark data. Finally, we specify the conditions in which hubness reduction is expected to be beneficial.

## Results

### RNA-seq data is prone to hubness

In order to evaluate the magnitude of the hub phenomenon in sequencing data, we started with bulk RNA-seq, as we wanted to document the effect of the dropout on hubness and intrinsic dimensionality of the data. We used three bulk datasets from The Cancer Genome Atlas (TCGA) and ARCHS^4^ repositories and added zeros in order to simulate the dropout effect, either in a simple manner, distributing the excess zeros uniformly in the expression matrix, or using Splatter^23^ (see Methods). We used previously developed tools to quantify the magnitude of the hubness phenomenon,^24, 25^ and we also evaluated the asymmetry of the k-NN graph, which we show to be an informative measure of hubness (see Methods). We worked on the PCA-transformed data, changing the number of Principal Components (PCs) to increase or decrease the data dimensionality. The skewness of the in-degree distribution of the k-NN graph (k-skewness) and the asymmetry estimators tend to increase with the dimension before reaching a plateau, for all datasets (Figure 1A). The same increase is seen for the estimators retrieving the number of hubs (2k-estimator) and antihubs (antihub estimator) but without plateauing. Regarding the 2k-estimator, it peaks at intermediate dimensions before decreasing: this is due to the fact that the distribution is strongly skewed, with such a large number of antihubs, that this estimator becomes irrelevant at high dimensions. We tested two other hub estimators, the maximum in-degree and the number of hubs retrieved as the number of cells with an in-degree outside their bell-shaped distribution (see Methods). They behave similarly with respect to the dimension (Supplementary Figure 2A,B). From those observations, we concluded that there are hubs in RNA-seq data, which appear already at intermediate dimensions, namely 10 PCs.

**Figure 1.**
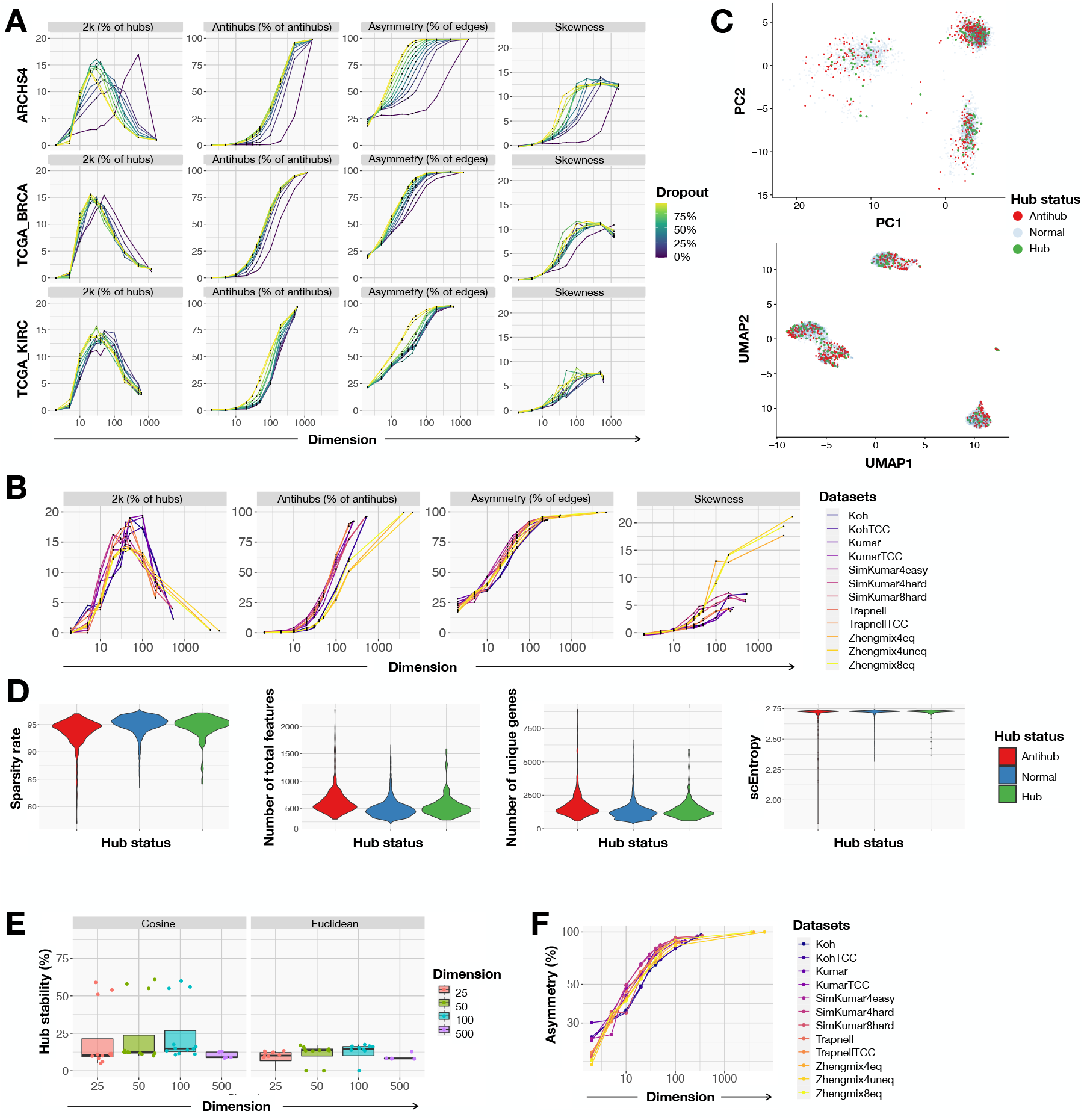
Hubness in sequencing data. We quantify hubness with 4 different estimators: percentage of hubs in the data defined as cells with an in-degree above 2*k*, using the same value for *k* as the one chosen to build the k-NN graph (first column), percentage of antihubs (second column), asymmetry of the k-NN graph (third column), skewness of the in-degree distribution (forth column). The quantification is shown as a function of the dimension after PCA reduction **(A**,**B)**. Hubness quantification methods are applied to 3 bulk datasets, with various rates of simulated dropout **(A)**, or to single-cell datasets **(B). (C)** Position of antihubs and reverse-coverage hubs in the PCA and UMAP projections; example with the Zhengmix4eq single-cell dataset. **(D)** Quality control metrics measured on antihubs, normal cells and hubs defined by reverse-coverage: dropout rate distribution (first column), number of total features counted (second column), number of unique genes detected (third column), single-cell entropy (scEntropy) distribution (forth column); example with the Zhengmix4eq single-cell dataset. **(E)** We count the proportion of reverse-coverage hubs that are in common between the original data and resampled data upon random removal of 10% of the cells, using scRNA-seq datasets.^26^ **(F)** Asymmetry of the k-NN graph over dimension upon removing of reverse-coverage hubs, with scRNA-seq datasets,^26^ in order to evaluate the resulting magnitude of the hubness phenomenon.

Concerning the effect of the dropout, it is negligible when the hub phenomenon is well established at the highest dimension, but it does lead to strong increase of hubness at intermediate dimensions (Figure 1A). This observation is reproducible across the two different methods used to generate dropout (Supplementary Figure 2C,D for the Splatter-simulated dropout). As the magnitude of the hub phenomenon reaches a plateau, we conclude that dropout drives the hubness up to this plateau.

To further investigate the link between sparsity and hubness, we studied their respective correlation with the global intrinsic dimension (GID, defined in Methods). Since an increase in data matrix sparsity expectedly involves a higher GID (R=0.93, p<0.0001, Spearman correlation) and a higher GID causes an increased hubness, defined with the asymmetry estimator, at k=10 and considering 100 dimensions (R=0.81, p<0.0001, Spearman correlation), we can assume that the effect of sparsity on hubness is at least partially due to the increased GID (Supplementary Figure 3A). This observation confirms also the intuition that hubness is a dimensional-related effect. We analyzed a diverse collection of scRNA-Seq datasets^26^ to measure the hub phenomenon. We used the same estimators as in Figure 1A and reproduced their evolution over increasing dimensionality (Figure 1B). We concluded that scRNA-seq data is prone to hubness as well, starting already from around 10 principal dimensions.

We also investigated the link between sparsity and hubness in scRNA-seq datasets, taking into account the GID value. In the case of single-cell data, we observed that sparsity is not enough to explain the variation of GID. We uncovered in fact three parameters that influence GID: the sparsity, the cardinality of the dataset, which corresponds to the number of observations, and the signal-to-noise ratio (SNR) (see Methods). The SNR and the cardinality are two dependent parameters (R=0.87, p=0.0026, Spearman correlation) (Supplementary Figure 3B), so we computed the correlation between GID and the composite parameter ratio of Sparsity to SNR, which appeared to be significant (R=0.77, p=0.015, Spearman correlation). Using the real single-cell datasets, we see that there is a weak correlation between the composite parameter Sparsity/SNR and the hubness (R=0.57, p=0.12, Spearman correlation), as well as between GID and hubness (R=0.29, p=0.45, Spearman correlation), using the asymmetry estimator with k=10 and 100 PCs to quantify hubness (Supplementary Figure 3C). Therefore, it seems that a higher Sparsity/SNR ratio is a partial explanation for a higher magnitude of the hub phenomenon, probably through its influence on GID.

### Identifying hubs in scRNA-seq data

Usual methods to retrieve hubs use a threshold on the hubness score, above which data points are defined as hubs. For example, the 2k-estimator counts those data points having their in-degree in the k-NN graph larger than 2k. In the case of single-cell transcriptomic datasets, large proportion of cells are antihubs, while the variance of the distribution of in-degree explodes: we mentioned that the tail follows a power law of Lévy, which means that whenever its slope in the log-log plot is smaller than -3, the variance diverges (Supplementary Figure 1B). As a consequence, the distribution of in-degrees is strongly skewed and the threshold-based methods fail, explaining the shape of the curve for the 2k-and the mean-estimators which do not reach a plateau (Figure 1A,B and Supplementary Figure 3A,B,D).

Instead, we suggest to directly use the neighborhood size in order to define the hub points. Considering the *N* cells with the highest in-degrees, one can calculate the proportion of total cells that have the latter *N* cells in their neighborhood lists, that we call “reverse-covered” cells. This quantity increases with *N*, reaching a plateau. We defined as hubs the *N* cells needed to hit the plateau, and called this method of hub identification “reverse coverage” (Supplementary Figure 4). In further sections, we use this method to retrieve hubs from scRNA-seq data.

### Hubs are not artefact cells

We assessed whether hubs have different properties compared to non-hub cells by looking at classical quality control (QC) metrics: number of genes, total number of features, dropout rate, entropy^27^ (see Methods), position in low-dimensional projections. We retrieved hubs using our reverse-coverage method.There was no clear difference in the distributions of the latter quality metrics between hubs, antihubs and other (normal) cells (Figure 1D and Supplementary Figure 5A). In some datasets, hubs and antihubs had more often a higher dropout rate, a higher number of unique genes detected or a lower number of total features compared to normal cells but this observation was not reproducible across all datasets; and there is no observable trend at all for the entropy. Regarding their position, the hubs and antihubs are scattered across the whole projection in the two-dimensional UMAP and PCA embeddings, in the sense that they do not form a distinct cluster or a set of outliers, although hubs seem to be located in denser regions (Figure 1C and Supplementary Figure 5B). In order to empirically rationalize the position of hubs we looked at the following model distributions in various dimensions: Gaussian and uniformly sampled hypercube. It was observed that for the Gaussian data, the distance to the data center decreased when the node degree increased except for the smallest dimensions which were less sensitive to hubness as expected. For the uniformly sampled hypercube data, in small dimensions the hubs were far from the origin (which was observed before^25^), nevertheless for high enough dimensions, hubs concentrated again near the data space origin (Supplementary Figure 6). Since the hubness phenomenon is specific to high dimensional data, we can conclude that hubs tend to concentrate near the cluster centers.

We also looked at the stability of hubs upon resampling of 90% of the cells and proved their poor stability: there is in most cases less than 25% of hubs in common between the original data and the resampled one, whatever the dimension and the metric used to compute the k-NN graph are (Figure 1E). It proves that being a hub is not an intrinsic property of the cells.

Lastly, we studied the hubness of the data after removing the hubs identified by the reverse-coverage approach, and observed that the k-NN graph remains asymmetrical, meaning that new hubs appeared in the data (Figure 1F). It also serves as a proof that hubness can not be reduced by merely removing hubs: more elaborated techniques are needed to correct the skewed k-NN graph.

Taken together, these observations suggest that hub or antihub cells do not form a distinct group in the data space: they are not explained by biological cell properties or technical artefacts. Hubs in single-cell datasets appear due to the effects of high-dimensional geometry and strongly impact the properties of cell neighbourhood graphs.

### Hubness reduction improves clustering accuracy

In order to study the effect of hubness reduction on the clustering of scRNA-seq data, we used quality-controlled datasets collected from previous clustering benchmark studies. We used those datasets which were specified to be of either gold-or silver-standard (see Methods, Supplementary Figure 7, Supplementary Table 1). For silver-standard datasets, one has to consider that label type and quality is a common limitation of studies assessing clustering accuracy. In particular, annotation of clusters with marker genes can differ as public marker databases (e.g. PanglaoDB and CellMarker) sometimes provide different indications.^28^ To compare these datasets in a uniform manner, we processed them using the Scanpy package^16^ according to standard steps with several combinations of parameters: quality control, normalization, log-transformation, gene selection, dimensionality reduction, scaling (Figure 2A).

**Figure 2.**
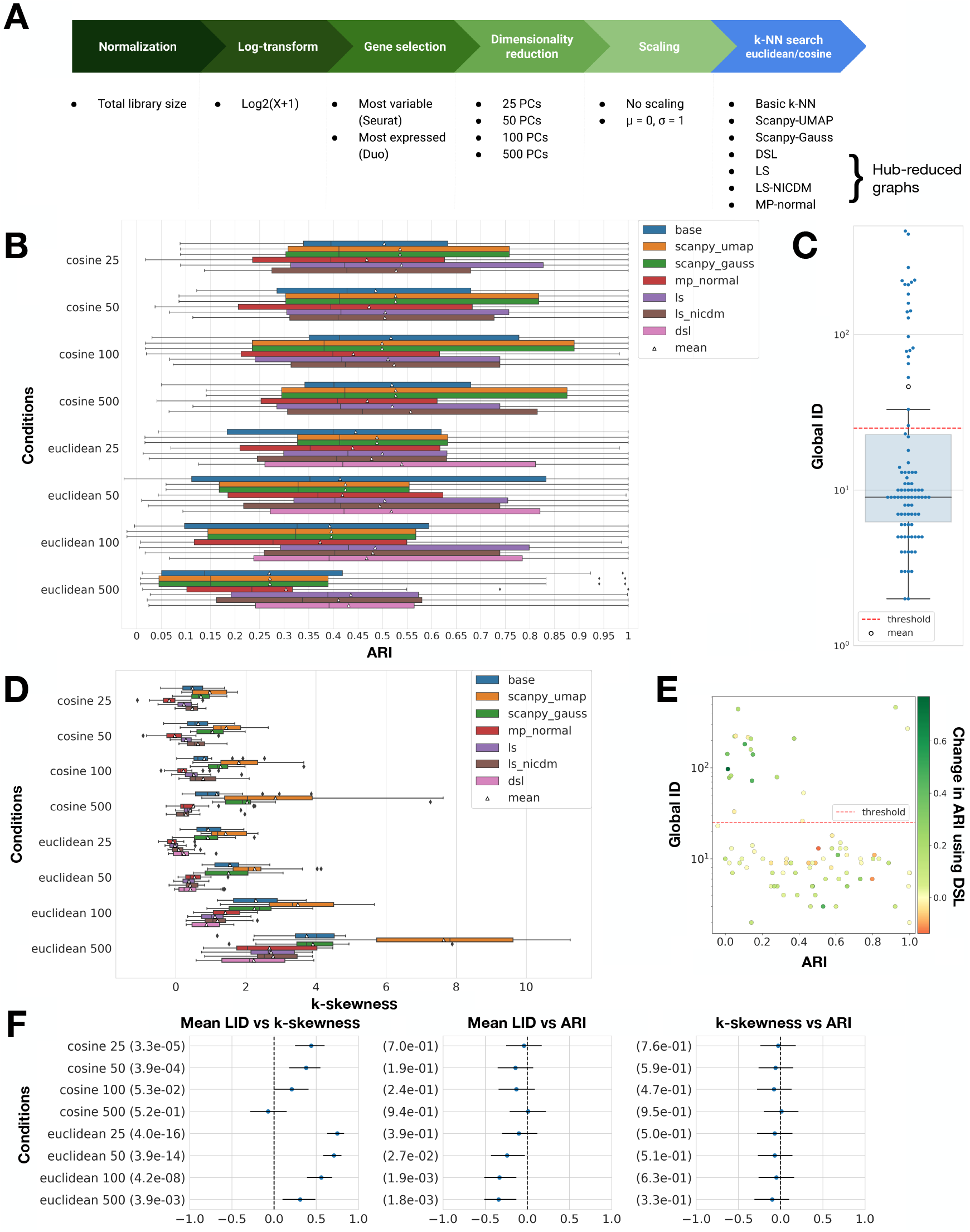
Evaluation of hubness reduction effect on clustering performance. **(A)** Preprocessing workflow with the different conditions used to construct various k-NN graphs upstream of the clustering task. **(B)** ARI scores for high-ID datasets, as a function of the metric, dimension and k-NN graph production method used; example with the Seurat recipe, scaling, and the Leiden algorithm. **(C)** Measure of the global ID (GID) of all datasets, used to define the high-ID datasets. **(D)** k-skewness of high-ID datasets, as a function of the metric, dimension and k-NN graph production method used; example with the Seurat recipe and scaling. **(E)** Relationship between global ID, ARI, and improvement in the clustering score using the hubness reduction algorithm DSL. **(F)** Pearson correlation coefficients distribution (p-value in parentheses) between k-skewness, ARI and mean local ID (LID) for all datasets, calculated with the base k-NN graph, using the Seurat recipe, scaling and the Leiden algorithm.

Hubness reduction was applied last, as a way to generate an alternative k-NN graph prior to clustering. We compared them with the uncorrected k-NN graph as well as with the k-NN graphs provided by two methods from the Scanpy package, based on UMAP and Gaussian kernel^21, 29^). Of note, the DSL hubness reduction is not compatible with the cosine dissimilarity. The clustering itself was performed using Leiden algorithm,^30^ which is widely used for single-cell data analysis as state-of-the-art clustering method. We also tested the Louvain algorithm^31^ which yielded similar results (corresponding figures are available on our Zenodo repository, DOI 10.5281/zenodo.4597151). Number of nearest-neighbors was set to the square root of the dataset cardinality (see Methods).

We followed previous studies and evaluated clustering accuracy with respect to ground truth labels using the Adjusted Rand Index (ARI) and the homogeneity scores.^32, 33^ Unlike the ARI, the homogeneity score does not penalize members of a class being split into several clusters and as such is complementary to the ARI. Since the Leiden method uses a resolution parameter rather than the desired number of clusters, and different number of clusters affect the clustering evaluation with ARI, we searched consistently for the resolution parameter yielding the same or closest possible number of clusters compared to ground truth labels^34^ (see Methods).

Our results show that GID and hubness are important parameters to consider when performing unsupervised clustering of single-cell datasets. We noticed that datasets with higher GID, i.e. above 25, had generally lower clustering scores evaluated with the ARI, whereas low and high scores were possible for lower GID (Figure 2E). The two exceptions of high-ID datasets with high clustering scores correspond to two simulated datasets included in our benchmark.^26^ These high-ID datasets are also the ones prone to hubness in the Euclidean space: indeed the mean local ID (LID) correlates with k-skewness. Although there is no direct correlation between k-skewness and ARI, it is therefore clear that hubness, as well as GID, need to be taken into account when performing clustering (Figure 2F).

Subsequently, hubness reduction was most useful for high-ID datasets (Figure 2C, Supplementary Figure 8), where clustering using hubness-reduced k-NN graphs performed better than using the base k-NN graph or the k-NN graphs from Scanpy (Figure 2B,E, Supplementary Figure 9). We also note that the use of cosine dissimilarity to build the k-NN graph resulted both in lower hubness and higher clustering accuracy than the Euclidean distance, as expected from previous literature^35^ (Figure 2D). Cosine dissimilarity’s sensitivity to hubness was also stable with regard to the number of PCs retained (Figure 2D). Interestingly, the highest average ARI and homogeneity scores were achieved using hubness reduction and 500 PCs. This provides a rationale to consider cosine dissimilarity and related metrics (e.g. the angular distance) as more robust and appropriate metrics to use for unsupervised clustering of single-cell data. It also indicates that a less stringent dimension reduction yield better clustering performance, which gives support to the hypothesis that drastic dimension reduction might lead to loosing useful information. For the case of low-ID datasets, the benefit of doing hubness reduction is not obvious anymore and should be tested on a per-dataset basis (Supplementary Figure 10).

To reflect the importance of taking into account how the ground truth was defined in these different datasets, we also looked at the power of hubness reduction considering only gold-standard datasets. Even if the improvement in ARI and homogeneity scores is less strong than if we consider only high-ID datasets, hubness reduction remains a useful step to perform, especially for the Euclidean distances, and considering a number of PCs above 25 (Supplementary Figure 11).

Dimension reduction and hubness reduction both mitigate negative effects of high GID on downstream analysis. Our study suggests that these two procedures have complementary effects, hubness reduction allowing to reduce the dimension less stringently. We also observe from Figure 3 that the improvement in clustering performance is accompanied by a more homogeneous density and a reduced skewness of the k-NN graph, while the k-NN graph constructed with the popular UMAP approach corrects only density inhomogeneity but not high skewness.

**Figure 3.**
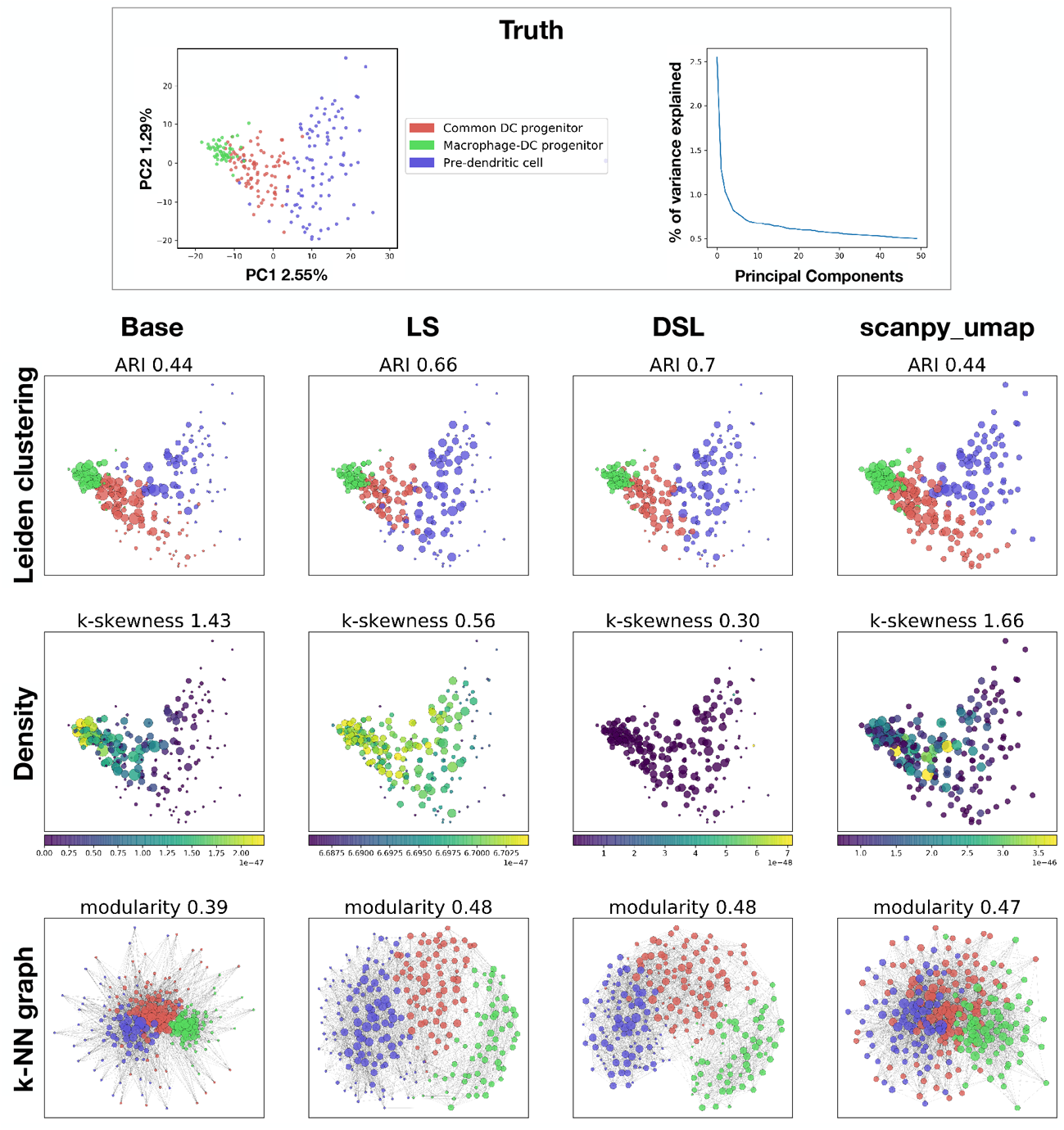
A selected example of Leiden clustering on a scRNA-seq dataset with FACS-labelled mouse blood dendritic cells (GSE60783), using Euclidean distance, 50 PCs and 15-NN graph. Clustering with the usual k-NN graph (base) or the UMAP k-NN graph (scanpy_umap) results in lower ARI while Local Scaling (ls) and DisSimLocal (dsl) k-NN graphs yield better accuracy. Both hubness-reduced and UMAP k-NN graphs produce more uniform Gaussian kernel density estimates. However, unlike hubness reduction, UMAP k-NN graph does not reduce the skewness of the in-degree distribution. The modularity is improved for the UMAP and hubness-reduced graphs compared to the base one, although the UMAP graph looks more intricate by eye. Point size is proportional to the in-degree in the respective k-NN graph.

To clarify this observation, we evaluated the strength of the density correction in a model high-dimensional Gaussian distribution which has been inhomogeneously sampled. Briefly, we take a Gaussian ball in 10 dimensions and remove 98% of the points in one of the half hyperball (Supplementary Figure 12A). As a mere consequence, the mean density of each half hyperball is different. We show that the density evaluated from the unweighted k-NN graph is more uniform after hub correction (Supplementary Figure 12C,D; see Methods). The local neighborhood relations are also better represented after the hubness reduction, in the sense that close points fall back in the same neighborhood (Supplementary Figure 12B).

As another evidence that hubness reduction improves clustering quality, we also tested this hypothesis using bulk RNA-seq datasets. We considered a collection of datasets from the ARCHS^4^ repository, constructed the k-NN graphs with or without hubness reduction, then ran Louvain algorithm and calculated the modularity of the resulting clustering. As one can see, hubness reduction improved modularity in the absolute majority of the datasets (Supplementary Figure 13). This analysis confirms that hubness reduction can be a useful step in many clustering pipelines.

### Hubness reduction improves trajectory inference in scRNA-seq datasets

In order to evaluate the effect of hubness reduction on the performance of trajectory inference (TI) in scRNA-seq data, we generated various k-NN graphs as input for the TI task, with or without hubness reduction. We used the Partition-based Graph Abstraction (PAGA)^36^ method for this purpose since it was ranked as the best TI tool.^37^ It is also very appropriate in our study since it is applied directly on k-NN graphs. We took the same parameters and preprocessing steps as for the clustering study described above (Figure 2A) except for the scaling step that we performed systematically in this experiment, to produce different trajectories that were then compared across the classical or hubness-reduced k-NN graphs using several quality metrics previously introduced.^37^ The following quality scores have been utilized: correlation to evaluate the relative position of cells along the trajectory, F1_branches to compare branch assignment and featureimp_wcor to measure the respective importance of differentially expressed features while constructing the trajectory (see Methods). We also calculate an overall score to average these three metrics. We tested our pipeline on the datasets from Sun et al. and the Cytotrace study,^38, 9^ first considering the expression matrices characterized by large values of GID (Supplementary Table 1).

We observed that the inferred trajectories were closer to the ground truth in most cases when TI was performed on a hub-reduced k-NN graph rather than using the base or the Scanpy k-NN graphs, in terms of the overall summary score and regardless of the combination of preprocessing parameters (Figure 4B). Exceptions were some combinations of preprocessing parameters used with 25 PCs (namely the Duo recipe with the cosine dissimilarity). Some combinations used with 100 PCs were not improved with hubness reduction either (namely the cosine dissimilarity with the Duo recipe and Leiden algorithm) (Supplementary Figures 16, 17). In total, only 9 out of 32 combinations of preprocessing parameters failed to yield better overall performance with hubness reduction. Out of these 9 combinations, 4 were computed with 25 PCs. When hub reduction is applied on the datasets embedded in lower dimensional spaces, e.g. 25 PCs, it is actually not surprising that hubness reduction has a weaker effect since the magnitude of hubness itself is smaller. Also, 8 combinations were computed with the cosine dissimilarity, which we know from previous experiments exhibit less initial hubness compared to the Euclidean metric (Figure 2D). To conclude, we noticed that the benefit of hubness reduction was much higher when using a reasonably high number of PCs and the Euclidean metric, which is coherent with the observations of the clustering task.

**Figure 4.**
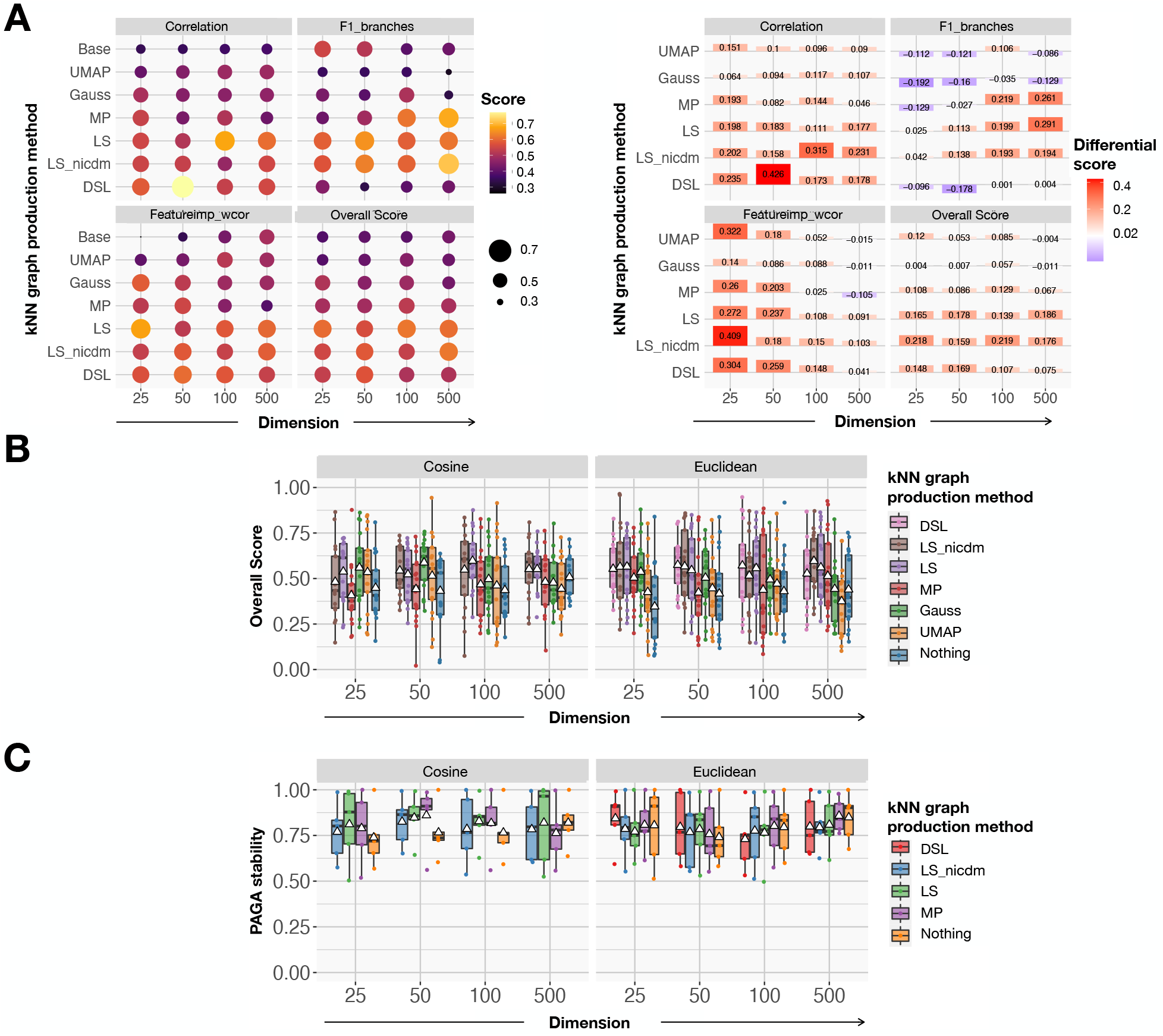
Trajectory inference (TI) improvement from application of hubness reduction. **(A)** Mean (left) and differential (right) TI quality scores (taking as reference the base score) for the three TI quality metrics and the summary score as a function of the dimension and the k-NN graph production method, calculated for the high-ID datasets;^38, 9^ example with the Seurat recipe, Euclidean metric and Leiden algorithm. **(B)** Overall detailed TI quality score of all datasets, as a function of the dimension and the k-NN graph production method, calculated for the high-ID datasets;^38, 9^ example with the Seurat recipe and Leiden algorithm. **(C)** Overall detailed PAGA stability score after subsampling 95% of the features and cells, as a function of the dimension and the hubness reduction method, calculated for the high-ID datasets;^38, 9^ example with the Seurat recipe and Leiden algorithm.

We display one example of preprocessing parameters combination (with the Seurat recipe, the Euclidean metric and the Leiden algorithm) in Figure 4A to show the improvement of the various TI quality scores compared to the base k-NN graph (Supplementary Figure 20). There are no clear patterns revealing that the increase in the quality of TI would be due to a specific increase in one of the three quality metrics: in fact, it depends highly on the preprocessing (Supplementary Figures 20, 21).

Briefly, we also noticed a slight partial improvement in the TI performance if we consider the low dimensional datasets from Sun et al.^38^ This is again especially true for the Euclidean metric, except for the preprocessing done with the Duo recipe and the Leiden algorithm (Supplementary Figures 14, 15, 18, 19). This is interesting compared to the clustering task, for which the dimensionality of the datasets was an important parameter to decide whether hubness reduction would be beneficial or not.

If we consider all the different preprocessing combinations and all datasets together, we can study the respective efficacy of each hubness reduction method. For the TI task and considering the highest dimensions where the magnitude of the hubness phenomenon is the strongest, we observed that the quality of the TI done after applying the two LS-based hubness reduction methods is the highest, shortly followed by DSL then MP. Going back to the datasets characterized by a low GID, it is not clear anymore what is the best hubness reduction method in order to improve TI. As a consequence, we suggest that one should test different hubness reductions, depending on their data, to reach the best performance, with a slight preference for the LS methods, especially when considering high dimensional datasets (Figure 4B).

Since it was mentioned^37^ that PAGA can be unstable, we followed their methodology to evaluate PAGA stability, using the base k-NN graph or the hubness-reduced ones. Again, we observed that the performance of PAGA, in terms of stability, increased in most cases whenever computed using the hubness-reduced graphs (Figure 4C). The benefit of using hubness reduction to improve stability depends on the method used. Surprisingly, while hubness reduction was most useful for the Euclidean distance when studying the mere performance, it turns out as most favorable to improve stability for the cosine dissimilarity. Furthermore, it is interesting to note that the efficacy of the different methods differs as well from the use case of pure TI performance. In the latter situation, MP was poorly performing, while it is quite efficient to improve PAGA stability.

### Low-dimensional embeddings upon hub reduction

To evaluate the impact of hubs and hubness reduction on the visualisation task, we designed two tests. Firstly, we used two distributions, the n-cube and the n-sphere, to evaluate the impact of hubness phenomenon on the goodness-of-fit between the projec-tion and the original data. The second test comprises scRNAseq data to quantify the quality of the projection before or after hubness reduction, in the same vein as what we did for the clustering and TI tasks. Here, we evaluated two visualisation algorithms, namely t-SNE^39^ and UMAP,^21^ that are widely used within the single-cell community.

We used randomly sampled n-cubes and n-spheres (see Supplementary Figure 22A), assuming that the n-cube exhibits hubs, while the n-sphere does not, or to a lesser extent (see Supplementary Figure 22B).^25^ Thus, we can estimate whether the presence of hubs impedes projecting the data onto a smaller (e.g. 2-dimensional) space. We quantify the goodness-of-fit by looking at the respective cost functions of the visualisation algorithms: Kullback-Leibler (KL) divergence and cross-entropy (CE) (see Methods), as well as at two metrics measuring correlation: the Quality of Distance Mapping (QDM) and the Quality of point Neighborhood Preservation (QNP; see Methods).^40^ The projection is the best possible whenever it minimizes the cost function and maximizes correlation. Our hypothesis is that the projection for a n-sphere will be of better quality than the one for a n-cube, because hubs distort the pairwise distance matrix used to compute t-SNE or UMAP. Regarding the cost functions, we note that they are designed to point towards the direction of the gradient descent for a given aim, but not as an absolute reference of the goodness-of-fit. We observed that QNP and QDM correlation metrics were always higher for the n-sphere than for the n-cube, both for t-SNE and UMAP, and irrespective of the number of dimensions or neighbors tested (see Figure 5A and Supplementary Figure 23). For the cost functions, and keeping in mind the fact that they focus on specific structures, we see that the KL divergence and the CE are smaller for the n-sphere than the n-cube, except for the CE computed after UMAP (see Figure 5B). We explain it by the fact that CE attributes a high importance to hubs and antihubs and thus the existence of these specific points accelerates the minimization of CE while performing the gradient descent. We reinforce this explanation with Figure 3, where we observe that the UMAP k-NN graph keeps the hubs at the center and the antihubs at the border, as in the base projection. Consequently, the k-NN graph structure with hubs is easier to preserve in the sense of the CE, even if the projection is overall of worse quality. Then we switched to single-cell datasets, using the same set of high-ID data as for the TI task, and tested the various k-NN graphs (the two Scanpy graphs and the four hub-reduced ones), but excluding the base one, that were projected in the UMAP, UMAP initialized with PAGA (PAGA+UMAP), or t-SNE spaces. This time, we evaluate the fit only with QNP and QDM metrics. To reduce the computation time, we evaluated less preprocessing combinations, using only the Seurat recipe, the Leiden clustering algorithm, scaling and 25, 50, 100 or 500 PCs. Looking at QDM and QNP in the different projected spaces, we see a reasonable improvement of the visualisation task performed after hub reduction, for at least one hub re-duction method, especially when using the Euclidean metric (see Figure 5C). There was only one use case for which the hubness reduction was not beneficial: when we projected the data with UMAP after PAGA initialization and with the cosine dissimilarity. For low-ID datasets, the benefit of using hubness reduction is not proven with our data (See Supplementary Figure 24).

**Figure 5.**
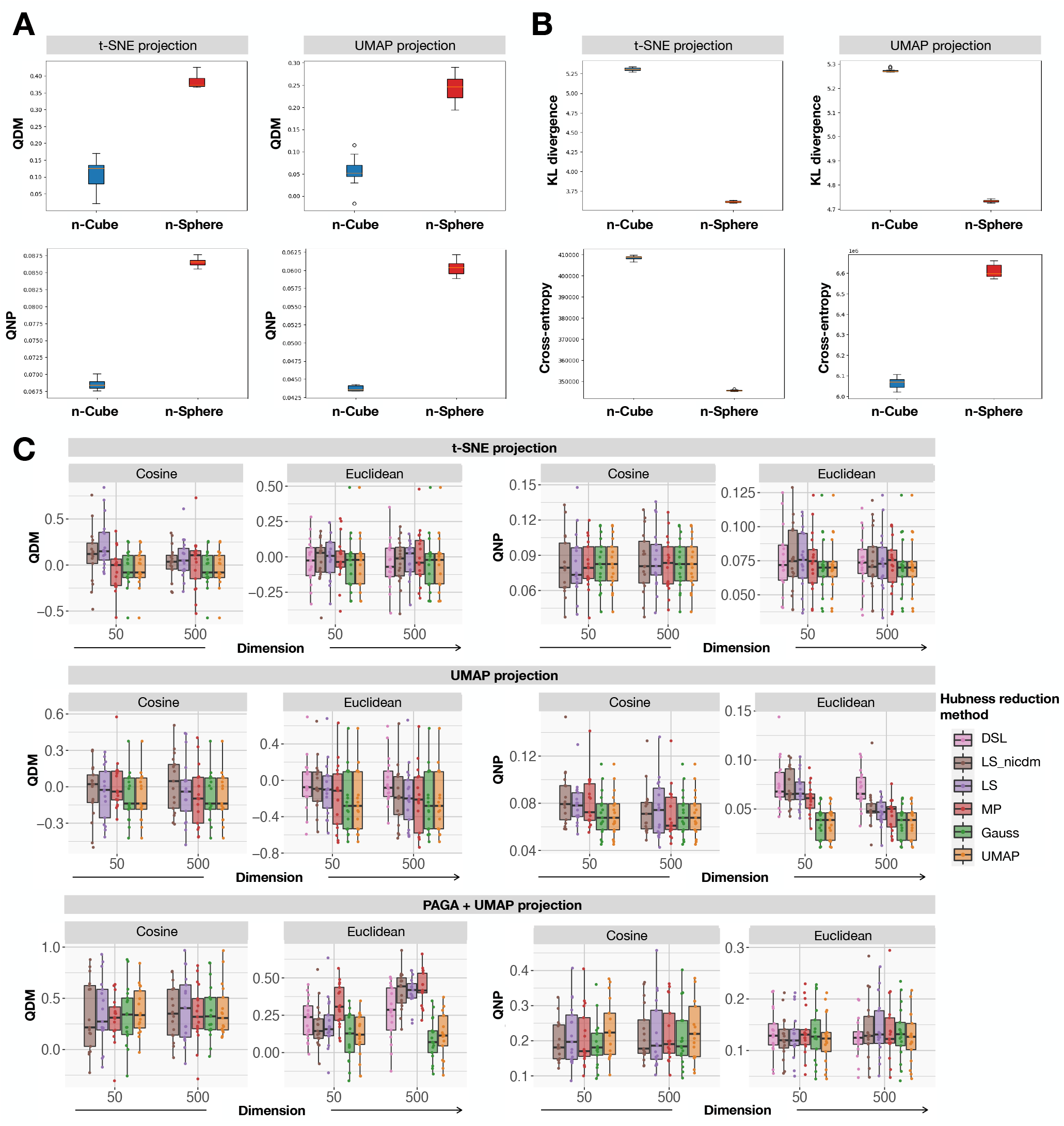
Effect of hubness reduction on the goodness-of-fit for t-SNE and UMAP. **(A)** QDM (top row) and QNP (bottom row) correlation metrics calculated on the t-SNE (left column) and UMAP (right column) projections of a 50-Cube and a 50-Sphere. The higher the correlation metric, the better the projection. **(B)** KL divergence (top row) and CE (bottom row) calculated on the t-SNE (left column) and UMAP (right column) projections of a 50-Cube and a 50-Sphere. The lower the cost function, the better the projection, in the sense of the quantity minimized by the cost function. **(C)** QDM and QNP before or after hubness reduction, evaluated after various visualisation algorithms, compared to the PCA with 500 PCs, for high-ID datasets. We project high-ID datasets either with t-SNE (top row), UMAP (middle row) or PAGA+UMAP (bottom row) and evaluate QDM (left column) and QNP (right column). The different projections are computed either with the cosine dissimilarity or the Euclidean metric, and using the two Scanpy k-NN graphs or the four hub-reduced graphs.

## Discussion

We proved that transcriptomic data is sensitive to the hubness phenomenon. Regarding the experiment with bulk data, we observed that all datasets were prone to hubness, but to various extent. We observed that this sensitivity positively correlates to the sparsity at least, most probably because the sparsity greatly influences the data GID. For the single-cell experiments, we showed the positive correlation between sparsity and the hubness phenomenon, even if this effect is mitigated by the cardinality and the signal-to-noise ratio. It would be very interesting to explore other reasons accounting for the difference in sensitivity to hubness.

In order to quantify the hubness phenomenon, we used methods that were previously introduced in the literature.^24,25^ They aim at counting either hubs or antihubs, or globally characterize the distribution of hubness scores across all data points. We concluded that for scRNA-seq data, methods of hubness quantification using threshold-based hubs counting do not perform adequately, since the hubness score distribution in high dimensions contains many values close to zero. Therefore we designed a method based on the size of the respective in-coming neighborhoods to retrieve hubs in a more robust way, that we called the reverse-coverage approach, that we used throughout our analyses.

We then studied the nature of these hubs, showing that they are not artefact cells or cells with specific biological properties. However, they have a topological utility, in the sense that they tend to be located close to the cluster centers and can be used for initialization of the clustering as such.^41^ One of the experiments to investigate the nature of hubs has been to remove them in the datasets and observe the persistence of the hubness phenomenon in the resampled data. Since the resampled data k-NN graph remains asymmetrical, it demonstrated that the simple hub removal is not a good approach for correcting the inhomogeneity of the k-NN graphs caused by high data dimensionality. In order to ameliorate the k-NN graph properties, we used instead existing techniques of hubness reduction modifying the local metric in the data space.^42, 24^

Regarding the clustering task, we show that hubness reduction can be beneficial, especially for the datasets characterized by high intrinsic dimensionality. As a matter of fact, clustering performs poorly on those datasets analysed with classical pipelines, probably because they suffer more from the unwanted hubness effects. Upon hub reduction, we are able to improve the clustering as seen with the increase of the standard clustering quality metrics ARI and homogeneity scores. We also noticed that cosine dissimilarity produces k-NN graphs that are less prone to the hubness phenomenon, compared to the more widely used Euclidean distance. To conclude, our results suggest that applying cosine dissimilarity and hub reduction can be beneficial for the task of clustering, especially for intrinsically high dimensional datasets. It indicates also that hub reduction is complementary to dimension reduction, while allowing to retain more principal components than is usually done. However, the available hub reduction methods differ in efficacy, with MP showing poor improvement in particular. We also extended this observation to bulk RNA-seq datasets. We empirically demonstrated that hubness reduction corrects for the presence of large density gradients and indegree connectivity distribution skewness in the k-NN graphs, outperforming other state-of-the-art methods such as the widely utilized UMAP algorithm, which might explain the better performances of the associated k-NN graphs for the clustering task.

Regarding the trajectory inference task, our results show that hubness reduction can improve the result of TI method application, as we exemplified using the popular PAGA algorithm. It is interesting to note that, while PAGA has already been shown as one of the best TI tools, it was done using the Euclidean metric and the UMAP or Gaussian approximate NN searches.^36^ In our case we improve further its performance with an alternative metric and NN search. Since PAGA works by pre-clustering the scRNA-seq data, we assume that the improvement in trajectory inference upon hubness reduction is at least partly due to the fact that the clustering task itself performs better. An interesting perspective would be to investigate thoroughly other leads of explanation for this improvement, and especially the differences between the different hub reduction methods. Our hypothesis to explain the poor performance of Mutual Proximity (MP) hub-ness reduction method for the clustering and TI tasks compared to the three other methods is that MP uses all pairwise distances to correct for hubness while we know that they suffer from the measure concentration. On the contrary, the other methods take advantage of the local neighborhoods which may explain their better efficiency. If we consider the impact of MP on k-skewness and measure concentration, it seems that its correction is the strongest out of the 4 methods, probably leading to over-corrected data (Supplementary Figure 25). It appears also interesting to test other TI approaches that rely on the construction of k-NN graph or other types of graphs approximating the data.^43^

Lastly, we showed that hubness reduction could improve visualisation of scRNAseq datasets, as expected looking at the previous literature.^44^ Again, it is especially true for high-ID datasets, as they suffer more from the hub phenomenon, and with the Euclidean metric which is more sensitive to the dimensionality.

### Materials and Methods

### Datasets used in benchmark

The bulk datasets were downloaded from the ARCHS^4^ and TCGA website. For the ARCHS^4^ expression matrix, we sampled the first 2000 observations to reduce the size of the dataset. We then filtered out manually 330 samples that were single cell RNA-seq or not RNA-seq samples (e.g. lncRNA, siRNA, etc), retaining a total of 1670 observations. For the TCGA datasets, we took all observations for breast (BRCA) and renal (KIRC) carcinomas.

To evaluate clustering and TI, we gathered single-cell datasets with gold-and silver-standard labels used in previous studies and benchmarks of supervised and unsupervised clustering and trajectory inference.^26,38, 28,45,46,9,37^ Labels on gold-standard datasets reflect ground truth information such as physical time points, genetic perturbations, cell lines, culture conditions, or FACS sorting. On the contrary, silver-standard datasets were labelled using a combination of FACS and unsupervised clustering.

### Hubness quantification

#### Hubness of simple model data distribution

We generated in Python Gaussian and uniformly sampled from hypercube data distributions with 10,000 samples, in spaces of dimension 2, 10, 50 and 500. Then we compute the 10-NN graph to retrieve the in-degree of each point, and show the distribution of in-degrees with 200 bins, averaged over 100 i.i.d. iterations for each dimensionality value and the data distribution.

#### Hubness scores of individual data points

From the bulk or single-cell datasets, we performed PCA using log-transformed data, while retaining only the 10,000 most variable genes:

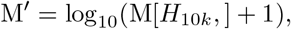

where *H* _10*k*_are the highly variable genes, M is the original gene-by-cell matrix and M’ the input matrix for the PCA.

Then the k-NN graph was computed using the RANN R library, with the number of PCs used ranging from 2 to the number of cells minus 1. For the value of k, we tested k ranging from 5 to 100, and displayed the results for k=10. From the k-NN graph, the in-degree or hubness score *X*_*i*_ is calculated for each cell *i*.

#### Hubness quantification for datasets

Let *k* be the value used to build the k-NN graph, *X* the distribution of hubness scores for individual data points, *µ* its mean and *σ*its standard deviation.

The 2*k* estimator counts the number of hubs, retrieved as the cells that have a hub score above 2*k*.

The *Mean* estimator counts the number of hubs, retrieved as the cells that have a hub sore above *µ* + 3 *σ*.

The *Antihub* estimator is the number of cells having zero hubness score.

The *Asymmetry* estimator counts the percentage of unidirectional edges in the k-NN graph.

The *Skewness* estimator is calculated as follows:

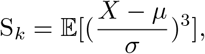

The *Maximum* estimator is the maximum hubness score observed in the distribution, divided by the cardinality of the dataset.

#### Dropout simulation

We simulated dropout in two different ways, using a Bernoulli distribution to decide which counts to drop. In the first setting, the dropout rate is a fixed constant for all samples, and we consider only non-zero gene counts. In the second setting, we used the tool from the R library Splatter to add dropout in a more realistic way, in order to reproduce the distribution of single-cell data.^23^ Briefly, the dropout rate of a given gene depends on its expression level following a logistic function.

#### Signal-to-Noise-Ratio (SNR) evaluation

To quantify the SNR, we assumed that the distribution of the eigenvalues from the cell-cell covariance matrix follows a Marcenko-Pastur distribution, except for a few eigenvalues that contain the signal of the data. As a consequence, we derive that the fraction of eigenvalues following the Marcenko-Pastur distribution is a good estimation of the noise magnitude, while the fraction of eigenvalues outside this distribution is a proxy for the signal magnitude. Since fitting the Marcenko-Pastur is not obvious, we designed a more simple proxy for SNR, that proved to be satisfactory in our experiments:

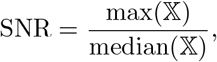

where 𝕏 is the distribution of the eigenvalues of the cell-cell covariance matrix.

#### Intrinsic dimensionality (ID) of datasets

We evaluated ID using PCA. Global ID is defined as the number of eigenvalues of the covariance matrix exceeding the first eigenvalue divided by 10 (so-called conditional number-based PCA ID estimate). We consider that datasets with a GID above 25 are high dimensional (high-ID datasets). Mean local ID is defined as the mean of ID values computed for the 100-nearest neighborhood of each point.

#### Reverse coverage method for identifying hubs

We suggest a novel definition of hubs based on the size of the in-coming neighborhood. Given a number *N* and taking the *N* cells with the highest hubness scores, we can calculate the number of cells *n* that have at least one of these *N* putative hubs in their nearest neighbors. We call these *n* cells the reverse-covered cells. Looking at the proportion of reverse-covered cells as a function of *N*, it reaches a plateau, meaning that the reverse coverage will increase only by a small fraction after the plateau. We define as hubs the *N* cells with the highest hubness scores, such that *N* is the first value for which we closely approach the plateau.

#### QC measurements

We used the Seurat library to compute the UMAP and PCA projections, as well as the total number of features and the number of unique genes. We computed in R the sparsity rate as the percentage of zeros in the expression matrix. To estimate the transcriptomic single-cell entropy, we used the scEntropy tool implemented in Python.^27^

To compute the stability of hub identity, we sampled 10 times 90% of the cells and looked at the mean percentage of hubs from the original data that were recovered in the resampled data to evaluate the intrinsic nature of hubs.

#### Hub positions with respect to the center of model data distributions

To evaluate hubs’ position, we generated in Python Gaussian and uniformly sampled in hypercube distributions with 105 points each, in different spaces of dimension 2, 5, 8, 10 and 20. We used the scikit-hubness library to construct the 10-NN graph and get the hubness score. From those scores we computed the average distance to the coordinate origin and its rank for the data points with a hubness score above a given threshold. To make the average computation more robust, we considered only the averages calculated over more than 100 points. For the average distance *M*_*t*_, we normalize it by the dimension:

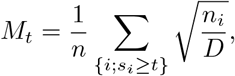

where *t* is the threshold on the hubness score, *D* the dimension, *s*_*i*_ the hubness score of point *i* and *n*_*i*_ its norm.

### Hubness reduction

#### Hubness reduction methods

We used the Python package scikit-hubness to measure skewness and to reduce hubness.^22^ This package offers 4 methods for reducing hubness, that produce a hub-corrected k-NN graph: Mutual Proximity (MP), Local Scaling (LS) and its variant LS-NICDM (Non-Iterative Contextual Dissimilarity Measure) and DisSimLocal (DSL).^42, 24^

Mutual Proximity models pairwise distances *d*_*i,j2{*1,…,*n}\i*_ of a set of *n* points with random variables *X*_*i*_ that depict the distribution of distances between *x*_*i*_ and all other points, then:

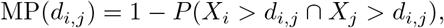

where *P* is the joint probability density function. Local Scaling is calculated using the pairwise distance *d*_*i,j*_ and takes into account the local neighborhood:

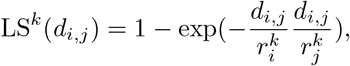

where *k* refers to the size of the local neighborhood, and 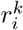 is the distance of point *x*_*i*_ to its k-th neighbor. The variant LS-NICDM uses the average distance to the *k* neighbors instead of the mere distance to the *k*-th neighbor:

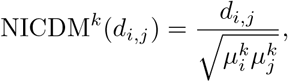

where 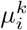 is the average distance of point *x*_*i*_ to its *k* nearest neighbors.

DisSimLocal uses local centroids *c*^*k*^(•) to reduce hubness:

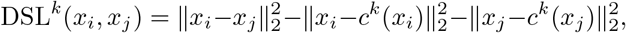

where the local centroid is estimated as the barycenter of the *k* nearest neighbors of *x*_*i*_:

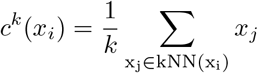

### Hubness and measure concentration

To evaluate the impact of hubness reduction on k-skewness and measure concentration in a general case, we generated two types of distributions: one or two Gaussian blobs in 10, 50 and 100 dimensions, with 5,000 points per blob, over 10 iterations. We used the scikit-hubness Python package to reduce hubness and measure k-skewness. For the measure concentration, we evaluated it as:

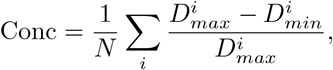

where 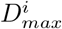 and 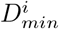are the maximum, resp. the minimum pairwise distances for point *i*.These distances were calculated either considering all points, or only the 50 nearest neighbors.

### Clustering

#### Dataset preprocessing

We processed the datasets used for the clustering task using the Python package Scanpy.^16^ We combined two recipes for the preprocessing (Duo or Seurat) with two different metrics (Euclidean and cosine dissimilarity), scaling or not, and four values for the number of PCs to retain (25, 50, 100, and 500, that we used to truncate the ambient data dimensionality in order to reduce the computational time). The following k-NN graphs were computed: simple (base) k-NN graph, four hub-reduced graphs, using the hub reduction methods provided by the Python scikit-hubness package, and two graphs provided by the Scanpy package with use of UMAP and Gaussian kernel methods^21, 29^). For the TI task, we used the same preprocessing pipeline, except for the scaling step, that we systematically perform.

The Duo recipe consists in log-normalizing the data, keeping the 5,000 most variable genes and normalizing again.

The Seurat recipe log-normalize the data as well and select the variable genes according to a set of thresholds: variable genes with a mean above 0.0125 and below 3, and a dispersion above 0.5. The data is normalized again after the gene filtering step.

The clustering was done on the seven k-NN graphs with the Leiden algorithm.^30^ The number of nearest-neighbors was set to the square root of dataset cardinality. Since the graph-based clustering methods do not allow choosing the exact number of clusters, we tuned the resolution parameter in order to get the ground truth number of clusters. We started with a resolution of 1.5 and limited the search of the resolution to the interval [0, 3]. Then we performed the clustering iteratively, with a maximal number of steps set to 20 and a resolution which would increase or decrease in a dichotomous manner:

##### Algorithm 1

Resolution tuning

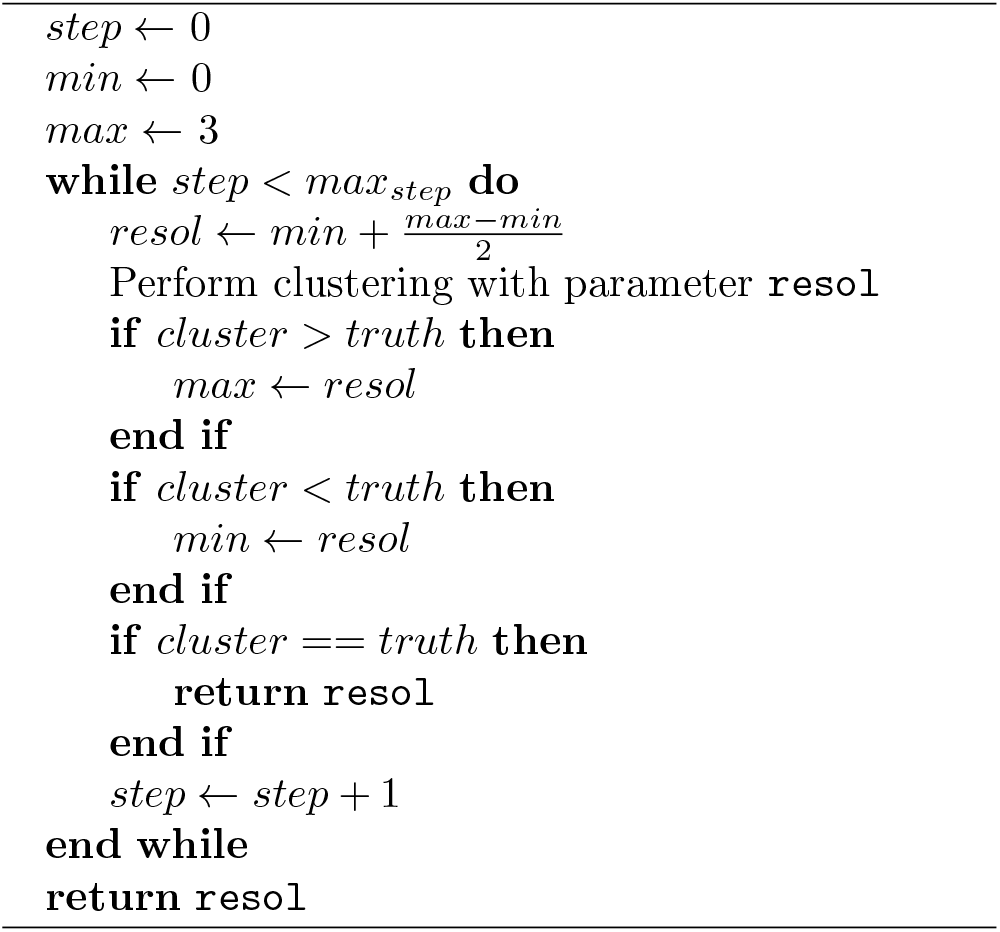

#### Evaluation of clustering accuracy

We used the Adjusted Rand Index (ARI) and the homogeneity scores to evaluate the quality of clustering.^33^ The best score is 1 and the worse is 0 for both measures.

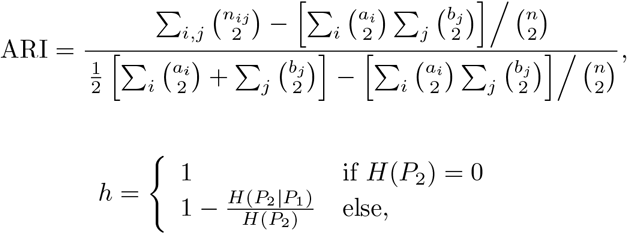

where

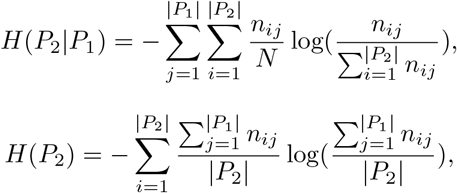

where *P*_1_ and *P*_2_ are the two partitions, *n*_*ij*_ is the value of the *i*-th row and *j*-th column in the contingency table, and *a*_*i*_, resp. *b*_*j*_, is the sum of the values sitting on the *i*-th row, resp. *j*-th column, of the contingency table.

#### Model distribution simulating strongly heterogeneous data point density

We generated a 10-dimensional Gaussian distribution containing 10,000 points. The data point cloud was separated in two parts by a hyperplane of coordinates x=0, x being the first axis. From the right half hyperball, we pick randomly 100 points and discard the others. We then constructed the k-NN graphs with the scikit-hubness Python package, with or without hubness reduction and used the k-NN graph to estimate the density.^47^ Briefly, it is possible to evaluate the following quantity from the unweighted k-NN graph:

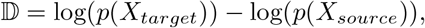

where *X*_*source*_ and *X*_*target*_ are two data points, and *p* is the local density.

First, we determine the shortest path, *y* between *X*_*source*_ and *X*_*target*_ in the k-NN graph using the Dijkstra algorithm. Then for each intermediate point *X*_*i*_ in the path, we get from the k-NN graph the quantities:

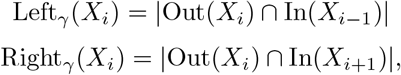

where In(*X*_*i*_) and Out(*X*_*i*_) are the in-and out-neighborhoods of *X*_*i*_. From this point, the density estimate along the path, *y* is:

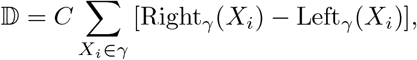

where *C* is a constant depending on *k* and the number of dimensions.^48^ In our case we fixed the source at the center of the Gaussian hyperball and randomly sampled 60 targets in each half hyperball, to be able to compare the estimates from the two half hyperballs by calculating the average of the density estimates for both half hyperballs.

#### Clustering modularity evaluation in bulk RNA-seq data

We took the mouse collection of datasets from the ARCHS^4^ data repository, retaining only those containing more than 300 samples. Without any other filters, it represents a total of 148 datasets.

To compute the modularity for each dataset, the data was log-transformed then projected or not in the PCA space with 50 components. From that we compute the k-NN graph with the cosine dissimilarity, with or without hub reduction done with the LS and MP methods only to reduce computation time. Finally we applied Louvain clustering algorithm using different k-NN graphs and computed the modularity *Q* using the Python library igraph:

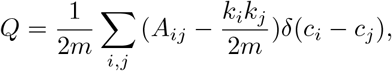

where *m* is the number of edges in the k-NN graph, *A*_*ij*_ is the element of the adjacency matrix on the *i*-th row and *j*-th column, *k*_*i*_ is the in-degree of point *i, c*_*i*_ its cluster identity and *δ* the Dirac function.

### Trajectory inference

#### Using PAGA algorithm

We used the implementation of PAGA from the Python package Scanpy. Same combinations of preprocessing steps, metrics and clustering algorithms have been used as described above in the clustering section.

### Evaluation of trajectories

We used the R toolbox dynverse to compute three quality metrics on each PAGA trajectory: correlation, F1_branches and featureimp_wcor.^37^

Correlation is calculated from the geodesic distance and quantifies the correlation between the relative distances of a given cell in the reference and the predicted trajectories:

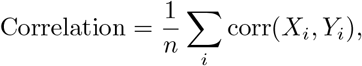

where *X*_*i*_ is the distribution of relative geodesic distances to cell *i* in the reference trajectory and *Y*_*i*_ the distribution of relative geodesic distances to *i* in the prediction.

F1_branches computes the similarity of branch membership between two trajectories, by mapping each cell to its closest branch:

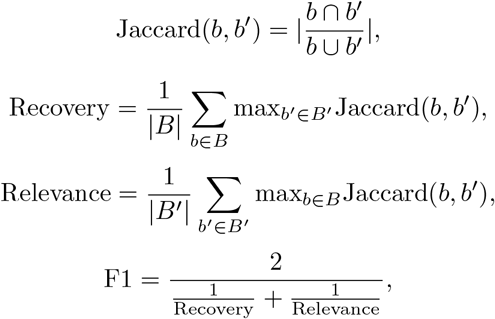

where B and B’ are the two branch partitions for the reference and the predicted trajectories.

For the calculation of featureimp_wcor, the geodesic distances of all cells to all milestones in the trajectory are computed, then predicted with a Random Forest.

From the Random Forest, we retrieve the importance of each gene for the prediction in the two trajectories in order to compute a weighted Pearson correlation, with the weights depending on the mean importance in the reference trajectory:

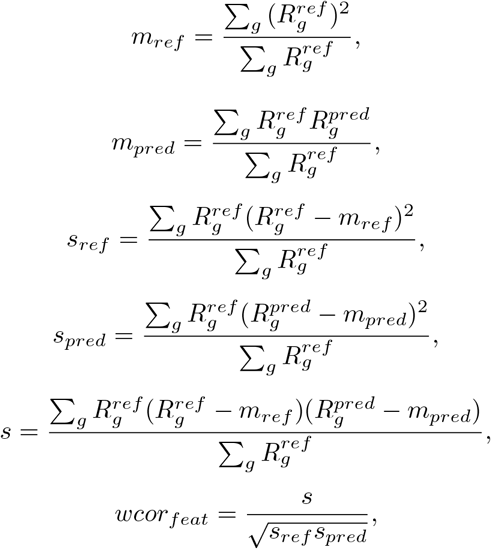

where *R*_*g*_ is the importance of gene *g* in the predicted or reference trajectory, *m* the weighted mean of the reference or the prediction, *s* the weighted variance and *s* the weighted covariance.

Each score was then normalized according to the following procedure: the scores of each metric were normalized and scaled such that *δ*=1 and *µ* =0 for each dataset, then we apply the unit probability density function of the normal distribution, to shift scores back in the [0,1] range. The calculation of a summary value for each metric merging all datasets for each preprocessing condition was performed by computing the arithmetic mean for the datasets with the same trajectory type and source, then the arithmetic mean fixing only the trajectory, and finally the arithmetic mean for all trajectory types. We also computed an overall score of the three quality metrics which is the arithmetic mean of the latter, either considering all the datasets or directly the different preprocessing.^37^

#### Trajectory stability

We used the same methodology described in a previous benchmark^37^ to evaluate the stability of PAGA. Briefly, we sample 95% of the cells and genes iteratively and evaluate the differences between two successive trajectories, doing 10 iterations and using the correlation and F1_branches metrics, but excluding featureimp_wcor which is not stable on the identity. To compare two successive iterations we compute both metrics using the common cells and genes, on the *n*+1 iteration, using the *n*-th iteration as the reference. We get a stability score for each metric. To compare the stability across the different datasets and conditions, we normalize the scores, such that the correlation and F1_branches have the same magnitude for each dataset. Briefly, for each dataset and each metric, we transform the scores to get *a-*=1 and *µ* = 1, then apply the unit probability density function of the normal distribution. We then compute the arithmetic mean of the two metrics. To speed the computation of the stability, we just ran it on the Sun et al. datasets^38^ and we did not compute the two scanpy k-NN graphs.

### Visualisation task

#### Generating n-cubes and n-spheres

We generated n-cubes and n-spheres in Python, using the packages scikit-dimension and numpy. We generated 10 sets for each distribution, each containing 5,000 points, embedded in spaces of various dimensions in the range [10, 50, 100].

#### Low dimension projections: t-SNE, UMAP, PAGA+UMAP

We used the following Python libraries to compute the projections: sklearn for t-SNE, umap for UMAP, and scanpy for PAGA+UMAP. For t-SNE, we use `metric=‘precomputed’` and `perplexity=50.0` for the single-cell experiment (and the default values for the model experiment). For PAGA+UMAP, we set `init_pos=‘paga’` when running `scanpy.tl.umap`. For all projections, we set `n_components=2`.

#### Correlation metric QDM and QNP

Quality of Distance Mapping quantifies the correlation of pairwise distances, only retaining a subset of the latter. We compute first what is called “natural PCA”^40^ on the reference: the pair of most distant points (*i*_1_, *j*_1_) represents the first components. Then, for the n+1 component (*i*_*n*+1_, *j*_*n*+1_), it is such that *i*_*n*+1_ is the most distant to the set of previous components *S*_*n*_ ={ *i*_1_,. .., *i*_*n*_, *j*_1_,. .., *j*_*n*_ } and *j*_*n*+1_ is the point of *S*_*n*_ closest to *i*_*n*+1_. We used this set of pair to compute the QDM:

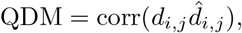

where *d*_*i,j*_ is the distance in the reference space, 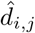 in the projection and we compute the correlation using the set of components *S*_*n*_ from the natural PCA. We took n=1000 for the tests with the hypercube and the hypersphere, and n equals to the number of cells for the tests with the single-cell datasets.

Quality of point Neighborhood Preservation computes the intersection of the neighborhoods in the reference and projection:

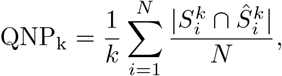

Where 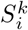, resp. 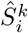, is the neighborhood of point i in the reference, resp. the projection, *k* is the size of the neighborhood and *N* the number of points. For the hypercube and hypersphere test, we took *k* in the range [10, 50, 100] and for the single-cell data, *k* equals the square root of the cardinality.

#### Cost function: Kullback-Leibler divergence and cross-entropy

The KL divergence is the cost function used for the t-SNE algorithm and is defined as:

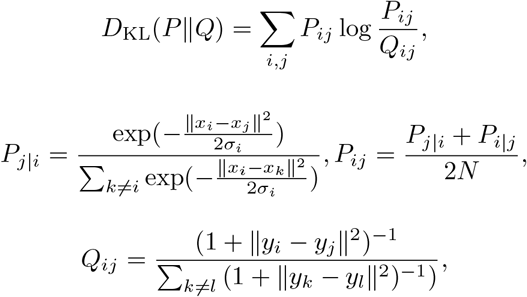

where *x*_*i*_, resp. *y*_*i*_, is the vector of the *i*-th point in the reference, resp. the projected, space, *a-*_*i*_ is a parameter that is entirely determined by the choice of the perplexity in the t-SNE algorithm and *N* is the number of points.

The cross-entropy is the cost function in UMAP:

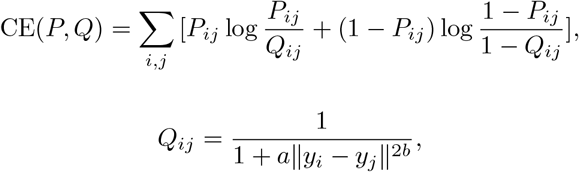

where *y*_*i*_ is the vector of the *i*-th point in the projected space, *a* and *b* are two parameters entirely determined by the choice of a *min*_*dist* in UMAP and *P*_*ij*_ is the membership strength of the 1-simplex between the *i*-th and the *j*-th points.

## Data and code availability

Data is available via Zenodo^49^ under DOI 10.5281/zenodo.4597151. Code has been uploaded on GitHub on the schubness repository.

## Acknowledgments

This work has been partially supported by the French government under management of Agence Nationale de la Recherche as part of the “Investissements d’Avenir” program, reference ANR-19-P3IA-0001 (PRAIRIE 3IA Institute), by the Ministry of Science and Higher Education of Russian Federation (Project No. 075-15-2020-808), by the Association Science et Technologie, the Institut de Recherches Internationales Servier and the doctoral school Frontières de l’Innovation en Recherche et Education-Programme Bettencourt. We thank also the ABiMS platform for access to their computing resources.

