## Supplementary Files for "Hubness reduction improves clustering and trajectory inference in single-cell transcriptomic data"

### Supplementary Tables and Figures

| Dataset name in benchmark study | Sequencing protocol | Number of cells | Number of features | Number of cluster labels | Description | Ref. | Benchmark study | Label type | Used to evaluate |
|---|---|---|---|---|---|---|---|---|---|
| Koh | SMARTer | 531 | 48981 | 9 | FACS purified H7 human embryonic stem cells in different differention stages | GSE85066 | Duo et al. PMC6134335 | FACS | Clustering |
| KohTCC | SMARTer | 531 | 811938 | 9 | FACS purified H7 human embryonic stem cells in different differention stages | GSE85066 | Duo et al. PMC6134335 | FACS | Clustering |
| Kumar | SMARTer | 246 | 45159 | 3 | Mouse embryonic stem cells, cultured with different inhibition factors | GSE60749 | Duo et al. PMC6134335 | Culture conditions | Clustering |
| KumarTCC | SMARTer | 263 | 803405 | 3 | Mouse embryonic stem cells, cultured with different inhibition factors | GSE60749 | Duo et al. PMC6134335 | Culture conditions | Clustering |
| SimKumar4easy | Synthetic dataset | 500 | 43606 | 4 | Simulation using different proportions of differentially expressed genes | PMC6134335 | Duo et al. PMC6134335 | Simulated data | Clustering |
| SimKumar4hard | Synthetic dataset | 499 | 43638 | 4 | Simulation using different proportions of differentially expressed genes | PMC6134335 | Duo et al. PMC6134335 | Simulated data | Clustering |
| SimKumar8hard | Synthetic dataset | 499 | 43601 | 8 | Simulation using different proportions of differentially expressed genes | PMC6134335 | Duo et al. PMC6134335 | Simulated data | Clustering |
| Trapnell | SMARTer | 222 | 41111 | 3 | Human skeletal muscle myoblast cells, differention induced by low-serum medium | GSE52529 | Duo et al. PMC6134335 | Culture conditions | Clustering |
| TrapnellTCC | SMARTer | 227 | 684953 | 3 | Human skeletal muscle myoblast cells, differention induced by low-serum medium | GSE52529 | Duo et al. PMC6134335 | Culture conditions | Clustering |
| Zhengmix4eq | 10x | 3994 | 15568 | 4 | Mixtures of FACS purified peripheral blood mononuclear cells | SRP073767 | Duo et al. PMC6134335 | FACS | Clustering |
| Zhengmix4uneq | 10x | 6498 | 16443 | 4 | Mixtures of FACS purified peripheral blood mononuclear cells | SRP073767 | Duo et al. PMC6134335 | FACS | Clustering |
| Zhengmix8eq | 10x | 3994 | 15716 | 8 | Mixtures of FACS purified peripheral blood mononuclear cells | SRP073767 | Duo et al. PMC6134335 | FACS | Clustering |
| GSE59114 | Smart-seq2 | 1622 | 7539 | 3 | Mouse Blood Phenotypes Aging HSCs (Smart-seq2) | GSE59114 | Gulati et al. PMID: 31974247 | FACS | Clustering |
| GSE74767 | SC3-seq | 212 | 28796 | 7 | Human (SC3-seq) | GSE74767 | Gulati et al. PMID: 31974247 | Cell lines | Clustering |
| GSE74767 | SC3-seq | 421 | 28796 | 13 | Macaque Embryo Timepoints Blastocyst timepoints (SC3-seq) | GSE74767 | Gulati et al. PMID: 31974247 | Timepoints | Clustering |
| GSE90860 | C1 | 223 | 42832 | 3 | Mouse Brain Timepoints Cortical interneurons (C1) | GSE90860 | Gulati et al. PMID: 31974247 | Timepoints | Clustering/TI/Visualisation |
| GSE95753 | 10x | 6000 | 27933 | 14 | Mouse Brain Phenotypes Dentate gyrus phenotypes (10x) | GSE95753 | Gulati et al. PMID: 31974247 | Markers (+clustering) | Clustering |
| GSE95753 | 10x | 6000 | 27933 | 8 | Mouse Brain Timepoints Dentate gyrus timepoints (10x) | GSE95753 | Gulati et al. PMID: 31974247 | Timepoints | Clustering |
| GSE67123 | Tang et al. | 143 | 24028 | 5 | Mouse Embryo Timepoints Embryonic HSCs (Tang et al.) | GSE67123 | Gulati et al. PMID: 31974247 | Timepoints | Clustering/TI/Visualisation |
| GSE98451 | CEL-seq | 714 | 12479 | 5 | Mouse Uterus Timepoints Endometrium (CEL-seq) | GSE98451 | Gulati et al. PMID: 31974247 | Timepoints | Clustering/TI/Visualisation |
| GSE99933 | Smart-seq2 | 369 | 23420 | 4 | Mouse Adrenal medulla Phenotypes Peripheral glia (Smart-seq2) | GSE99933 | Gulati et al. PMID: 31974247 | Markers (+clustering) | Clustering |
| GSE94641 | Plate-seq | 225 | 33327 | 4 | Mouse Brain Timepoints Medial ganglionic eminence (C1) | GSE94641 | Gulati et al. PMID: 31974247 | Timepoints | Clustering/TI/Visualisation |
| GSE60783 | C1 | 248 | 15752 | 3 | Mouse Blood Phenotypes Dendritic cells (C1) | GSE60783 | Gulati et al. PMID: 31974247 | FACS | Clustering/TI/Visualisation |
| GSE67602 | C1 | 1422 | 25932 | 5 | Mouse Skin Phenotypes Hair epidermis (C1) | GSE67602 | Gulati et al. PMID: 31974247 | Markers (+clustering) | Clustering/TI/Visualisation |
| GSE70245 | C1 | 394 | 23955 | 8 | Mouse Blood Phenotypes HSPCs (C1) | GSE70245 | Gulati et al. PMID: 31974247 | FACS | Clustering/TI/Visualisation |
| GSE90047 | Smart-seq2 | 447 | 40829 | 7 | Mouse Liver Timepoints Hepatoblast (Smart-seq2) | GSE90047 | Gulati et al. PMID: 31974247 | Timepoints | Clustering |
| GSE75748 | C1 | 1018 | 19095 | 6 | Human Embryo Phenotypes hESC in vitro (C1) | GSE75748 | Gulati et al. PMID: 31974247 | FACS | Clustering |
| GSE52529 | C1 | 170 | 46077 | 3 | Human Muscle Phenotypes HSMM (C1) | GSE52529 | Gulati et al. PMID: 31974247 | Culture conditions | Clustering/TI/Visualisation |
| GSE85066 | C1 | 498 | 30670 | 9 | Human Embryo Phenotypes Mesoderm (C1) | GSE85066 | Gulati et al. PMID: 31974247 | FACS | Clustering |
| GSE93421 | 10x | 5000 | 16957 | 10 | Human Blood Phenotypes Peripheral blood (10x) | GSE93421 | Gulati et al. PMID: 31974247 | Markers (+clustering) | Clustering |
| GSE36552 | Tang et al. | 85 | 20012 | 6 | Human Embryo Phenotypes Pre-implant human embryo (Tang et al.) | GSE36552 | Gulati et al. PMID: 31974247 | Timepoints | Clustering |
| GSE86146 | Smart-seq2 | 1844 | 24153 | 17 | Human Embryo Timepoints Germ cells (Smart-seq2) | GSE86146 | Gulati et al. PMID: 31974247 | Timepoints | Clustering |
| GSE98664 | RamDA-seq | 456 | 47515 | 5 | Mouse Embryo Timepoints mESC in vitro (RamDA-seq) | GSE98664 | Gulati et al. PMID: 31974247 | Timepoints | Clustering |
| GSE52583 | C1 | 101 | 23093 | 4 | Mouse Lung Timepoints Lung development (C1) | GSE52583 | Gulati et al. PMID: 31974247 | Timepoints | Clustering/TI/Visualisation |
| GSE97391 | inDrop | 2684 | 28205 | 4 | Mouse Brain Phenotypes Direct in vitro neuron (inDrop) | GSE97391 | Gulati et al. PMID: 31974247 | Markers (+clustering) | Clustering |
| GSE76408 | CEL-seq | 480 | 23460 | 6 | Mouse Intestine Phenotypes Lgr5-CreER intestine (CEL-seq) | GSE76408 | Gulati et al. PMID: 31974247 | Markers (+clustering) | Clustering |
| GSE109774 | 10x | 3652 | 13526 | 11 | Mouse Blood Phenotypes Bone marrow (10x) | GSE109774 | Gulati et al. PMID: 31974247 | FACS (+clustering) | Clustering |
| GSE109774 | Smart-seq2 | 4897 | 17479 | 8 | Mouse Blood Phenotypes Bone marrow (Smart-seq2) | GSE109774 | Gulati et al. PMID: 31974247 | FACS (+clustering) | Clustering |
| GSE92332 | Smart-seq2 | 1522 | 20108 | 9 | Mouse Intestine Phenotypes Intestine (Smart-seq2) | GSE92332 | Gulati et al. PMID: 31974247 | Markers (+clustering) | Clustering |
| GSE97391 | inDrop | 2996 | 28205 | 7 | Mouse Brain Phenotypes Standard in vitro neuron (inDrop) | GSE97391 | Gulati et al. PMID: 31974247 | Markers (+clustering) | Clustering |
| GSE45719 | Smart-seq2 | 286 | 22431 | 13 | Mouse Embryo Phenotypes Pre-implant mouse embryo (Deng et al.) | GSE45719 | Gulati et al. PMID: 31974247 | Timepoints | Clustering |
| GSE52583 | C1 | 66 | 23093 | 3 | Mouse Lung Phenotypes AT2/AT1 lineage (C1) | GSE52583 | Gulati et al. PMID: 31974247 | Markers (+clustering) | Clustering/TI/Visualisation |
| GSE69761 | C1 | 79 | 35016 | 5 | Mouse Lung Phenotypes Lung fibroblast (C1) | GSE69761 | Gulati et al. PMID: 31974247 | Markers (+clustering) | Clustering/TI/Visualisation |
| GSE92332 | Drop-seq | 4581 | 15971 | 15 | Mouse Intestine Phenotypes Intestine (Drop-seq) | GSE92332 | Gulati et al. PMID: 31974247 | Markers (+clustering) | Clustering |
| GSE107122 | Drop-seq | 5998 | 21201 | 3 | Mouse Brain Timepoints Neural stem cells (Drop-seq) | GSE107122 | Gulati et al. PMID: 31974247 | Timepoints | Clustering |
| GSE64447 | C1 | 447 | 24480 | 4 | Mouse Bone Phenotypes Skeletal stem cells (C1) | GSE64447 | Gulati et al. PMID: 31974247 | FACS | Clustering/TI/Visualisation |
| GSE102066 | C1 | 781 | 13762 | 8 | Human Brain Timepoints In vitro NPCs (C1) | GSE102066 | Gulati et al. PMID: 31974247 | Timepoints | Clustering |
| GSE75330 | C1 | 5050 | 23226 | 12 | Mouse Brain Phenotypes Oligodendrocyte phenotypes (C1) | GSE75330 | Gulati et al. PMID: 31974247 | Markers (+clustering) | Clustering |
| GSE75330 | C1 | 5050 | 23226 | 23 | Mouse Brain Timepoints Oligodendrocyte timepoints (C1) | GSE75330 | Gulati et al. PMID: 31974247 | Timepoints | Clustering |
| GSE87375 | Smart-seq2 | 338 | 40829 | 6 | Mouse Pancreas Timepoints Pancreatic alpha cell (Smart-seq2) | GSE87375 | Gulati et al. PMID: 31974247 | Timepoints | Clustering |
| GSE87375 | Smart-seq2 | 575 | 40829 | 7 | Mouse Pancreas Timepoints Pancreatic beta cell (Smart-seq2) | GSE87375 | Gulati et al. PMID: 31974247 | Timepoints | Clustering |
| GSE103633 | Drop-seq | 21612 | 28065 | 2 | Planaria Organism Phenotypes Whole planaria (Drop-seq) | GSE103633 | Gulati et al. PMID: 31974247 | Markers (+clustering) | Clustering |
| GSE107910 | Drop-seq | 9307 | 19530 | 8 | Mouse Thymus Timepoints Thymus (Drop-seq) | GSE107910 | Gulati et al. PMID: 31974247 | Timepoints | Clustering |
| GSE106587 | Drop-seq | 39505 | 23974 | 12 | Zebrafish Organism Timepoints Early zebrafish (Drop-seq) | GSE106587 | Gulati et al. PMID: 31974247 | Timepoints | Clustering |
| FreytagGold | 10x | 925 | 58302 | 3 | Human lung adenocarcinoma cell lines | GSE111108 | Freytag et al. PMC6124389, Sun et al. PMC6902413 | FACS | Clustering |
| PBMC3k | 10x | 3205 | 58302 | 11 | Human | SRP073767 | Freytag et al. PMC6124389, Sun et al. PMC6902413 | FACS | Clustering |
| PBMC4k | 10x | 4292 | 58302 | 11 | Human | SRP073767 | Freytag et al. PMC6124389, Sun et al. PMC6902413 | FACS | Clustering |
| Baron (Mouse) | inDrop | 1886 | 14861 | 13 | Mouse pancreas | GSE84133 | Abdelaal et al. PMC6734286 | Markers (+clustering) | Clustering |
| Baron (Human) | inDrop | 8569 | 17499 | 14 | Human pancreas | GSE84133 | Abdelaal et al. PMC6734286 | Markers (+clustering) | Clustering |
| Muraro | CEL-Seq2 | 2122 | 18915 | 9 | Human pancreas | GSE85241 | Abdelaal et al. PMC6734286 | FACS (+clustering) | Clustering |
| Segerstolpe | SMART-Seq2 | 2133 | 22757 | 13 | Human pancreas | E-MTAB-5061 | Abdelaal et al. PMC6734286 | Markers (+clustering) | Clustering |
| Xin | SMARTer | 1449 | 33889 | 4 | Human pancreas | GSE81608 | Abdelaal et al. PMC6734286 | Markers (+clustering) | Clustering |
| CellBench1 | 10X chromium | 3803 | 11778 | 5 | Mixture of five human lung cancer cell lines | GSE118767 | Abdelaal et al. PMC6734286 | Cell lines | Clustering |
| CellBench2 | CEL-Seq2 | 570 | 12627 | 5 | Mixture of five human lung cancer cell lines | GSE118767 | Abdelaal et al. PMC6734286 | Cell lines | Clustering |
| TM | SMART-Seq2 | 54865 | 19791 | 55 | Whole Mus musculus | GSE109774 | Abdelaal et al. PMC6734286 | FACS (+clustering) | Clustering |
| AMB | SMART-Seq v4 | 12832 | 42625 | 4/22/110 | Primary mouse visual cortex | GSE115746 | Abdelaal et al. PMC6734286 | FACS (+clustering) | Clustering |
| Zheng sorted | 10X CHROMIUM | 20000 | 21952 | 10 | FACS-sorted PBMC | SRP073767 | Abdelaal et al. PMC6734286 | FACS | Clustering |
| Zheng 68K | 10X CHROMIUM | 65943 | 20387 | 11 | PBMC | SRP073767 | Abdelaal et al. PMC6734286 | Markers (+clustering) | Clustering |
| Baron_m2016 | inDrop | 1886 | 14861 | 13 | Mouse pancreas | GSE84133 | Krzak et al. PMC6918801 | Markers (+clustering) | Clustering |
| Klein2015 | inDrop | 2712 | 24027 | 4 | Embryonic stem cells | GSE65525 | Krzak et al. PMC6918801 | Markers (+clustering) | Clustering |
| Zeisel2015 | STRT-Seq UMI | 3005 | 19972 | 9 | Mouse cortex and hippocampus | GSE60361 | Krzak et al. PMC6918801 | Markers (+clustering) | Clustering |
| Darmanis2015 | C1 | 466 | 21630 | 9 | Human brain | GSE67835 | Krzak et al. PMC6918801 | Markers (+clustering) | Clustering |
| Deng2014_raw | Smart-Seq, Smart-Seq2 | 268 | 21297 | 6 | Mouse embryo | GSE45719 | Krzak et al. PMC6918801 | Timepoints | Clustering |
| Goolam2016 | Smart-Seq2 | 124 | 28147 | 4 | Mouse embryo | E-MTAB-3321 | Krzak et al. PMC6918801 | Timepoints | Clustering |
| Kolodiejczyk2015 | SMARTer | 704 | 32225 | 3 | Mouse embryonic stem cells | E-MTAB-2600 | Krzak et al. PMC6918801 | Culture conditions | Clustering |
| Li2017 | SMARTer | 561 | 43055 | 9 | Human colorectal tumors | GSE81861 | Krzak et al. PMC6918801 | Markers (+clustering) | Clustering |
| Romanov2016 | C1 | 2881 | 21143 | 7 | Mouse hypotalamus | GSE74672 | Krzak et al. PMC6918801 | Markers (+clustering) | Clustering |
| Tasic2016_raw | SMARTer | 1679 | 21617 | 17 | Mouse cortical cells | GSE71585 | Krzak et al. PMC6918801 | FACS (+clustering) | Clustering |
| Deng2014_rpkm | Smart-Seq, Smart-Seq2 | 268 | 22958 | 5 | Mouse embryo | GSE45719 | Krzak et al. PMC6918801 | Timepoints | Clustering |
| Segerstolpe2016 | Smart-Seq2 | 3514 | 25525 | 15 | Human pancreatic islet cells | E-MTAB-5061 | Krzak et al. PMC6918801 | Markers (+clustering) | Clustering |
| Tasic2016_rpkm | SMARTer | 1679 | 24057 | 17 | Mouse cortical cells | GSE71585 | Krzak et al. PMC6918801 | FACS (+clustering) | Clustering |
| Yan2013 | Tang et al. | 90 | 20214 | 6 | Human embryo | GSE36552 | Krzak et al. PMC6918801 | Timepoints | Clustering |
| Biase2014 | SMARTer | 56 | 25737 | 4 | Mouse embryo | GSE57249 | Krzak et al. PMC6918801 | Markers (+clustering) | Clustering |
| Treutlein2014 | SMARTer | 80 | 23271 | 5 | Mouse lung epithelium | GSE52583 | Krzak et al. PMC6918801 | Markers (+clustering) | Clustering |
| ChuBatch1 | SMARTer | 350 | 19097 | 5 | Human | GSE75748 | Sun et al. PMC6902413 | FACS | Clustering/TI/Visualisation |
| ChuBatch2 | SMARTer | 425 | 19097 | 6 | Human | GSE75748 | Sun et al. PMC6902413 | FACS | Clustering/TI/Visualisation |
| Schlitzer | Fluidigm | 238 | 4480 | 3 | Mouse DCs from the BM | GSE60783 | Sun et al. PMC6902413 | FACS | TI/Visualisation |
| Petropoulos | Smart-Seq2 | 1289 | 8772 | 5 | Human embryo | E-MTAB-3929 | Sun et al. PMC6902413 | Timepoints | TI/Visualisation |
| LiM | Smart-Seq2 | 649 | 4777 | 8 | Male fetal gonads | GSE86146 | Sun et al. PMC6902413 | Timepoints | TI/Visualisation |
| LiF | Smart-Seq2 | 666 | 5319 | 12 | Female fetal gonads | GSE86146 | Sun et al. PMC6902413 | Timepoints | TI/Visualisation |
| ZhangBeta | Smart-Seq2 | 562 | 6138 | 7 | Mouse pancreatic beta cells | GSE87375 | Sun et al. PMC6902413 | Timepoints | TI/Visualisation |
| ZhangAlpha | Smart-Seq2 | 322 | 6138 | 6 | Mouse pancreatic alpha cells | GSE87375 | Sun et al. PMC6902413 | Timepoints | TI/Visualisation |
| GuoF | Tang et. al. | 100 | 8772 | 5 | Human female primordial germ cells | GSE63818 | Sun et al. PMC6902413 | Timepoints | TI/Visualisation |
| GuoM | Tang et. al. | 166 | 8772 | 5 | Human male primordial germ cells | GSE63818 | Sun et al. PMC6902413 | Timepoints | TI/Visualisation |
| KowalczykYoung | Smart-Seq | 493 | 2227 | 3 | Mouse stem cells | GSE59114 | Sun et al. PMC6902413 | FACS | TI/Visualisation |
| KowalczykOld | Smart-Seq | 873 | 2815 | 3 | Mouse stem cells | GSE59114 | Sun et al. PMC6902413 | FACS | TI/Visualisation |
| Hayashi | RamDA-seq | 414 | 23658 | 5 | Mouse ES cells | GSE98664 | Sun et al. PMC6902413 | Timepoints | TI/Visualisation |
| ShalekLPS | Smart-seq | 504 | 4158 | 5 | Mouse DCs | GSE48968 | Sun et al. PMC6902413 | Timepoints | TI/Visualisation |
| Trapnell | SMARTer | 290 | 8772 | 4 | Human skeletal muscle myoblasts cells | GSE52529 | Sun et al. PMC6902413 | Timepoints | TI/Visualisation |
| Olsson | SMARTer | 316 | 3594 | 3 | Mouse stem cells | GSE70245 | Sun et al. PMC6902413 | FACS | TI/Visualisation |

Supplementary Table 1: Table of all benchmark datasets' technical characteristics used in our study. Rows in gold are gold-standard, the rest are silver-standard datasets.

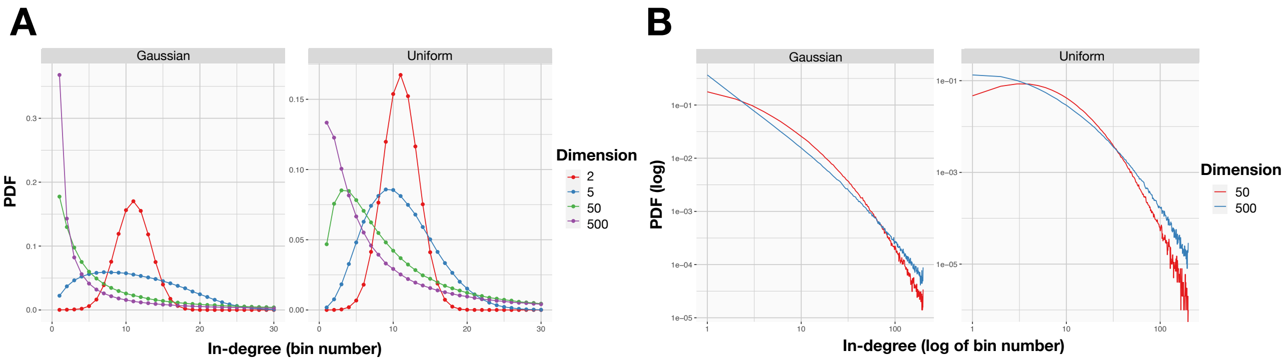


Supplementary Figure 1: Distribution of in-degree values. **(A)** Density of in-degrees for Gaussian (left panel) and uniformly sampled hypercube (right panel) distributions, with dimensions from 2 to 500; a fat tail appears with higher dimensions in both distributions. **(B)** Log-log plot of in-degree distributions for Gaussian (left panel) and uniformly sampled hypercube (right panel) distributions, with dimensions from 2 to 500; the fat tail can be linearly approximated, in both distributions.

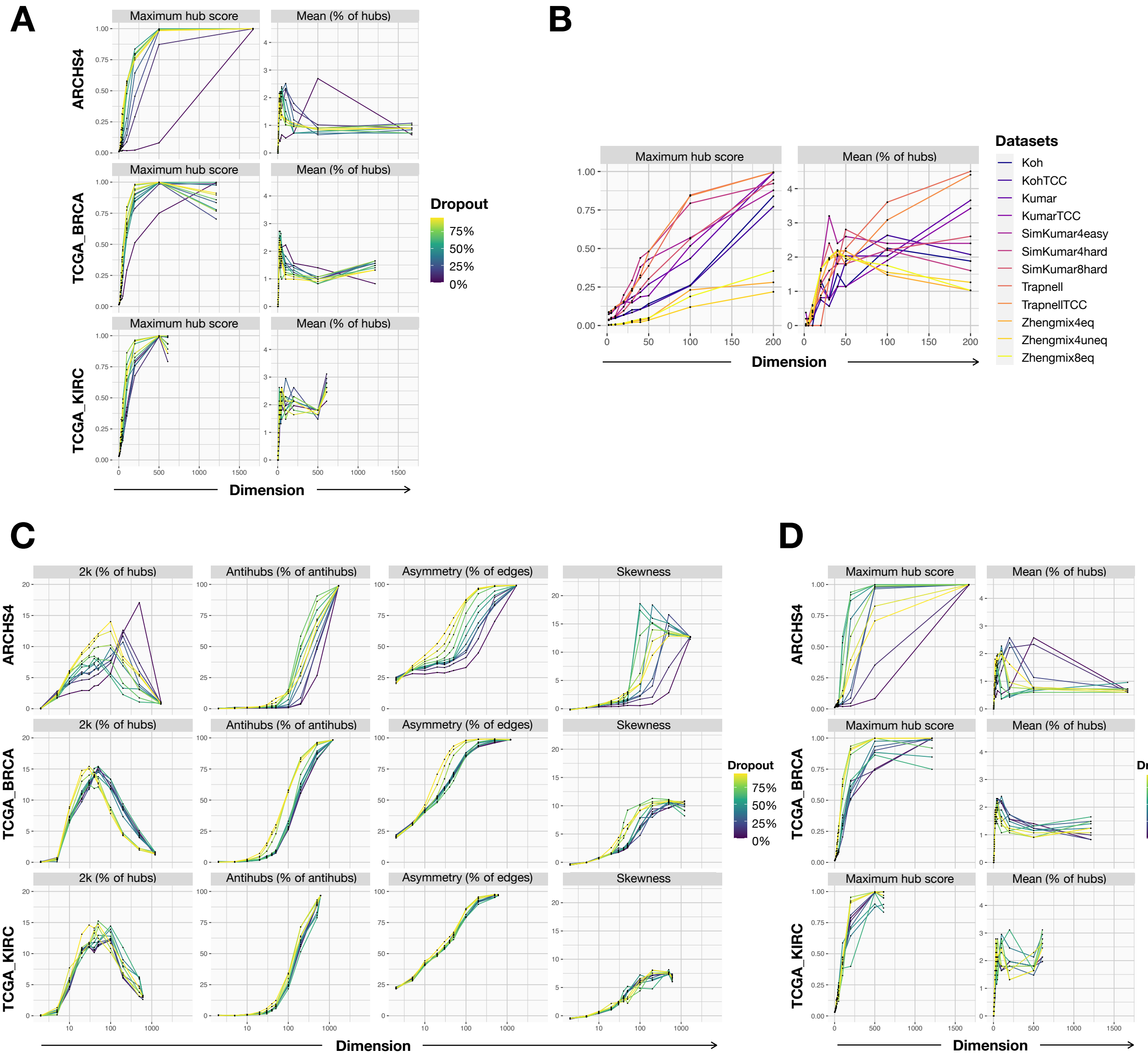


Supplementary Figure 2: Hubness in sequencing data. We quantify hubness with 2 alternative different estimators: maximum hubness score (first column), percentage of hubs as cells with an in-degree above $\mu+3\sigma$ (second column). The quantification is shown as a function of the dimension after PCA reduction **(A,B)**. Hubness quantification methods are applied to 3 bulk datasets, with various rates of simulated dropout **(A)**, or to single-cell datasets from[38] **(B)**. **(C)** Classical hubness quantification methods applied to bulk datasets, with various rates of Splatter-simulated dropout. **(D)** Alternative hubness quantification methods applied to bulk datasets, with various rates of Splatter-simulated dropout.

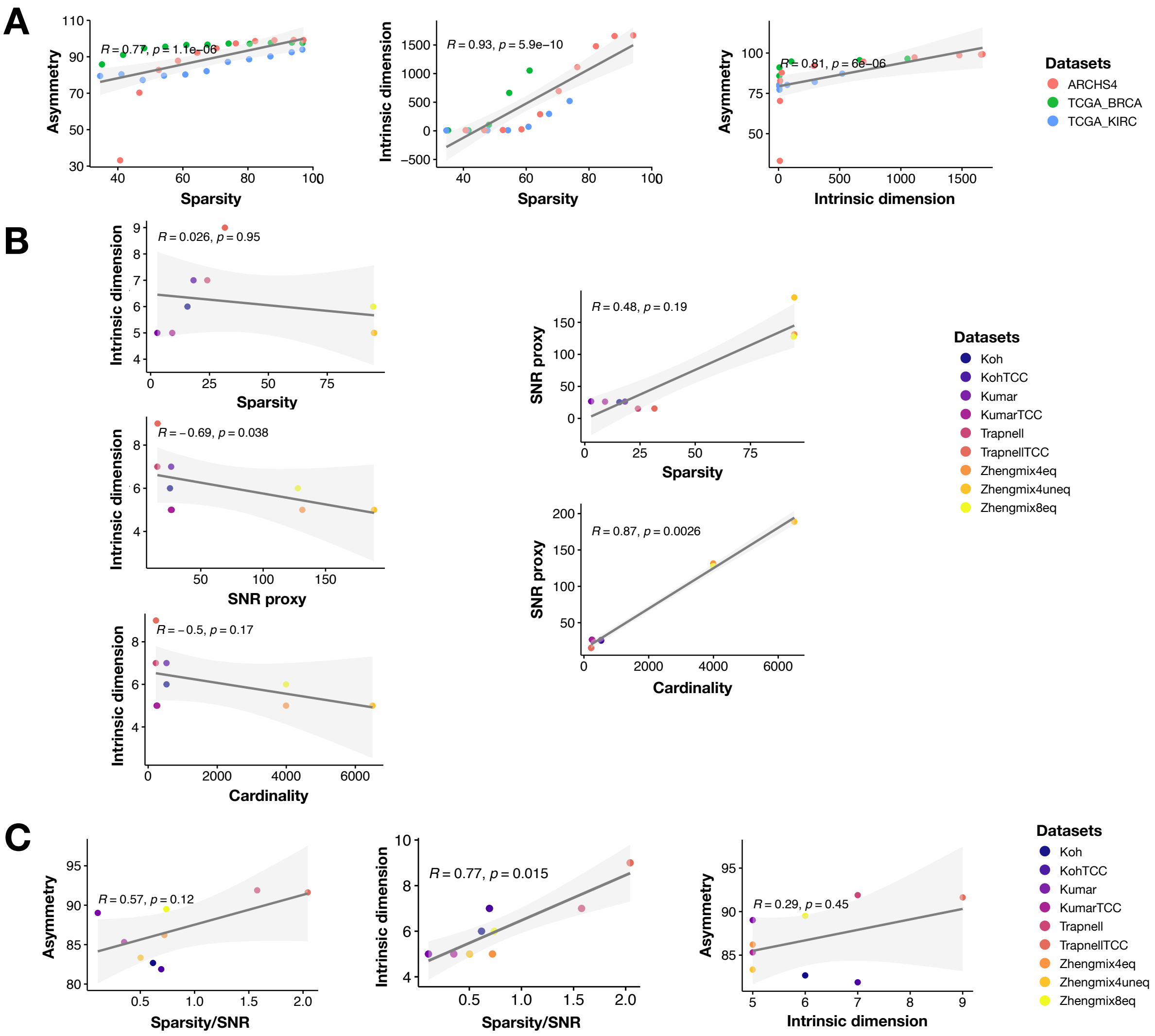


Supplementary Figure 3: Emergence of hubness relates to sparsity, SNR and GID. **(A)** We investigate the link between hubness and sparsity, by showing the Pearson correlation of sparsity to GID and of GID to hubness, using the bulk datasets with various rates of simulated dropout. **(B)** In the first column, we show the correlation between three parameters and GID: sparsity (first row), SNR (second row) and cardinality (third row). In the second column, we test the independence of these three parameters: SNR and sparsity are independent (first row) while SNR and cardinality are dependent (second row). **(C)** We investigate the link between hubness and the ratio of sparsity to SNR, by showing the Pearson correlation of the ratio sparsity/SNR to GID and of GID to hubness, using the 9 real single-cell datasets from.[38]

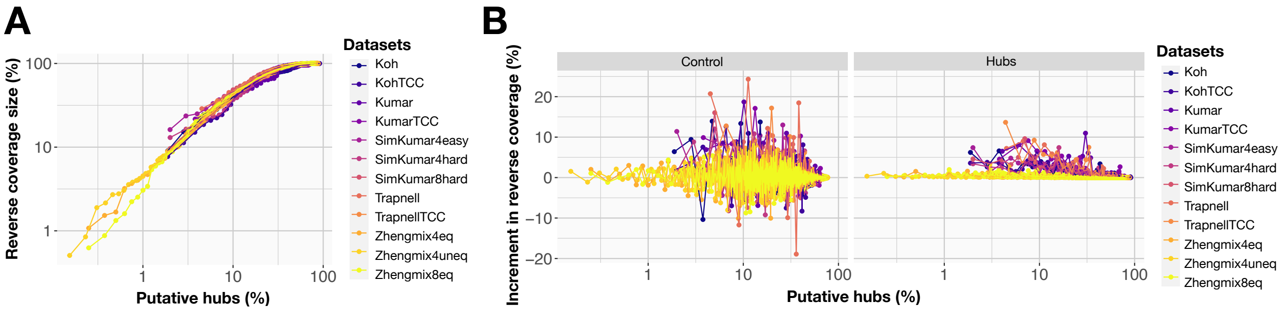


Supplementary Figure 4: Reverse-coverage hub-retrieving method. **(A)** Size of the reverse coverage (or percentage of reverse-covered data) as a function of the number of putative hubs, i.e. the number of cells with the highest in-degrees, done with the single-cell datasets from.[38] **(B)** Increment of the size of the reverse coverage, either as a function of $N$ random cells (left) or the $N$ cells with the highest in-degrees, considered as putative hubs (right), done on the same single-cell datasets.

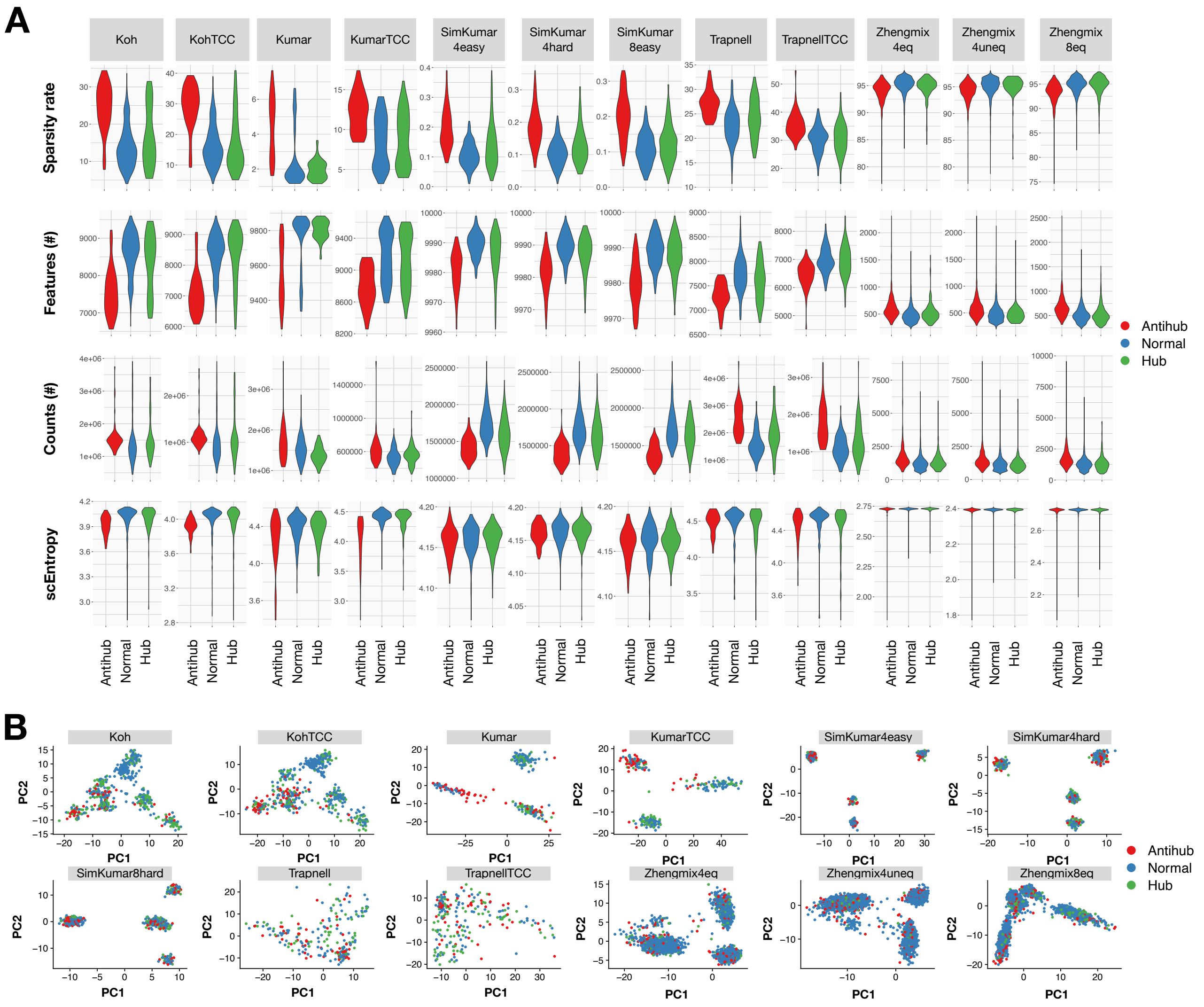


Supplementary Figure 5: Quality control metrics. **(A)** Quality control metrics distribution for hubs, antihubs and normal cells on the 12 datasets from Duo et al. (38): dropout rate (first row), number of total features (second row), number of unique genes (third row), single-cell entropy (last row). **(B)** PCA projections showing the positions of hubs, antihubs and normal cells for the 12 Duo datasets.

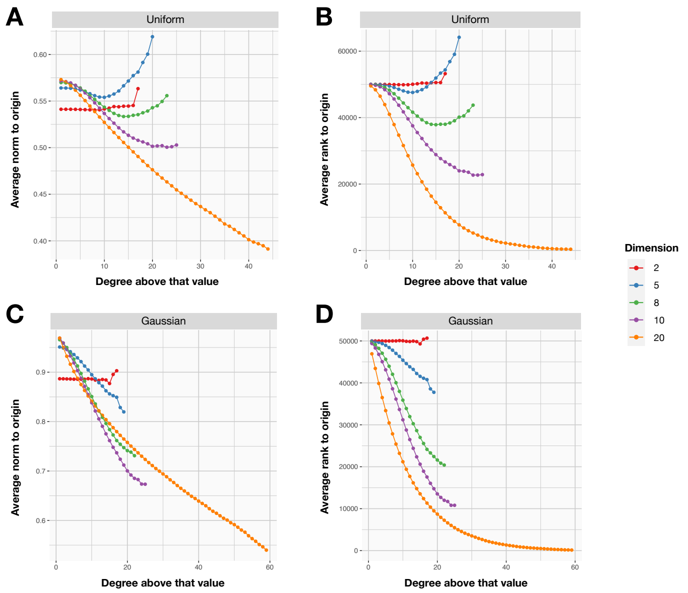


Supplementary Figure 6: Hubs positions. Position of hubs for uniform **(A,B)** and Gaussian **(C,D)** data distribution. **(A,C)** Average norm of points with an in-degree above the abscissa value. **(B,D)** Average ranking to the origin of points with an in-degree above the abscissa value.

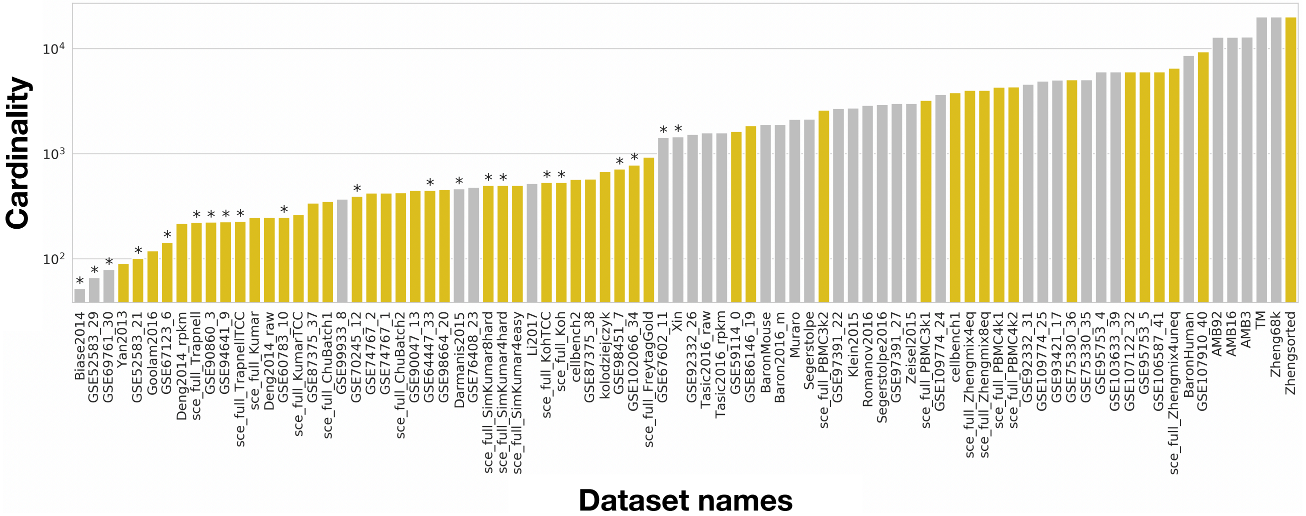


Supplementary Figure 7: Benchmark datasets collected from previous studies. Colors represent gold and silver standards, stars mark high ID. Note that the figure contains less datasets than listed in the supplementary table because there are overlaps between[9] and other benchmark studies

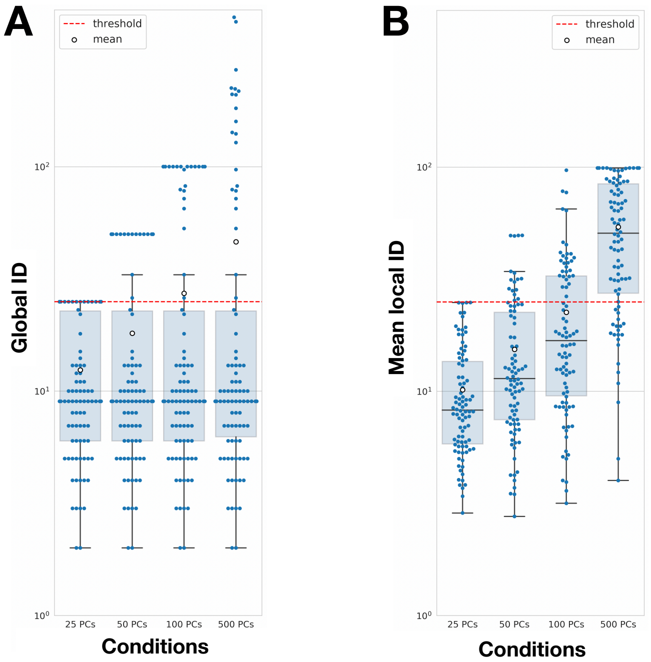


Supplementary Figure 8: GID **(A)** and mean LID **(B)** of datasets estimated with PCA, the Seurat preprocessing and scaling.

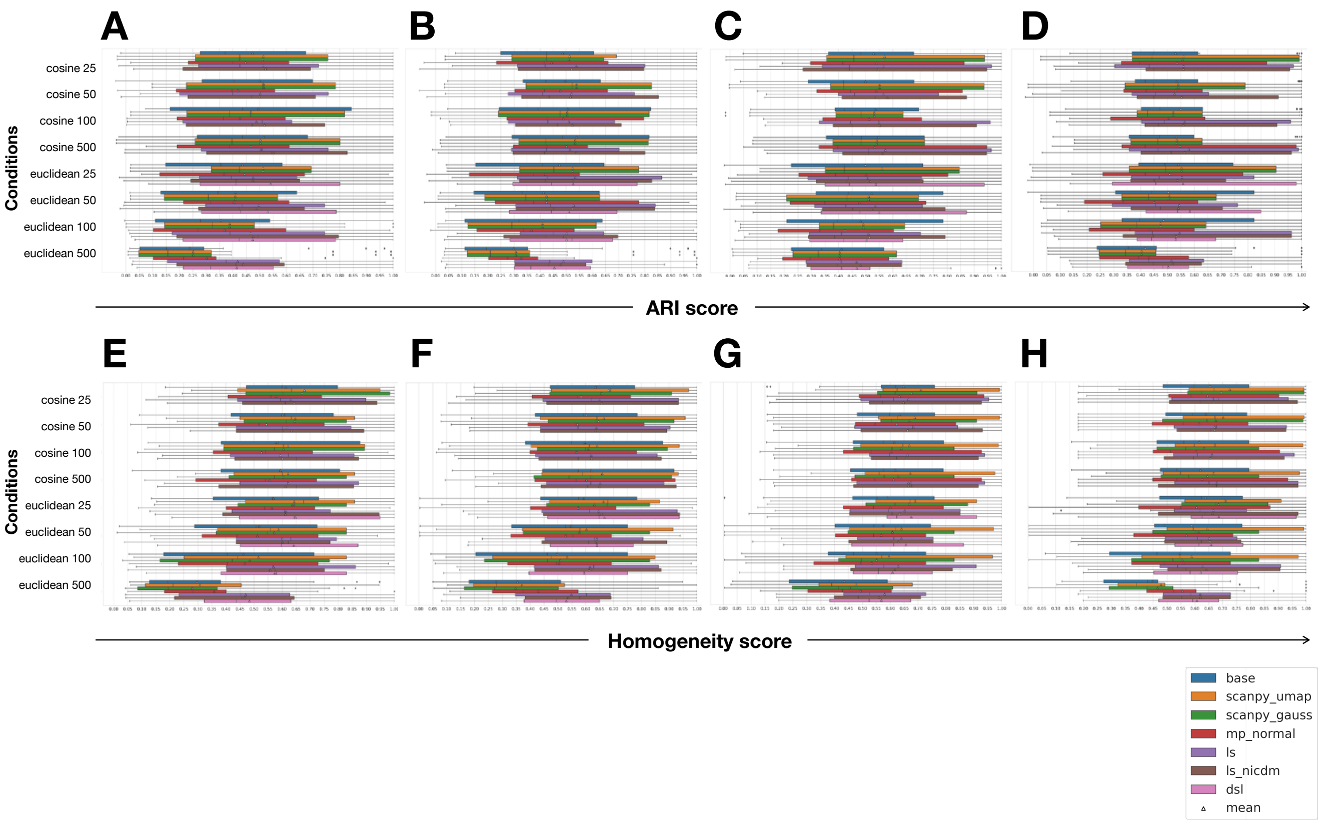


Supplementary Figure 9: clustering scores done with the Seurat **(A,B,E,F)** or Duo **(C,D,G,H)** preprocessing with **(A,C,E,G)** or without **(B,D,F,H)** scaling, for high-ID datasets: ARI scores **(A,B,C,D)** and homogeneity scores **(E,F,G,H)**.

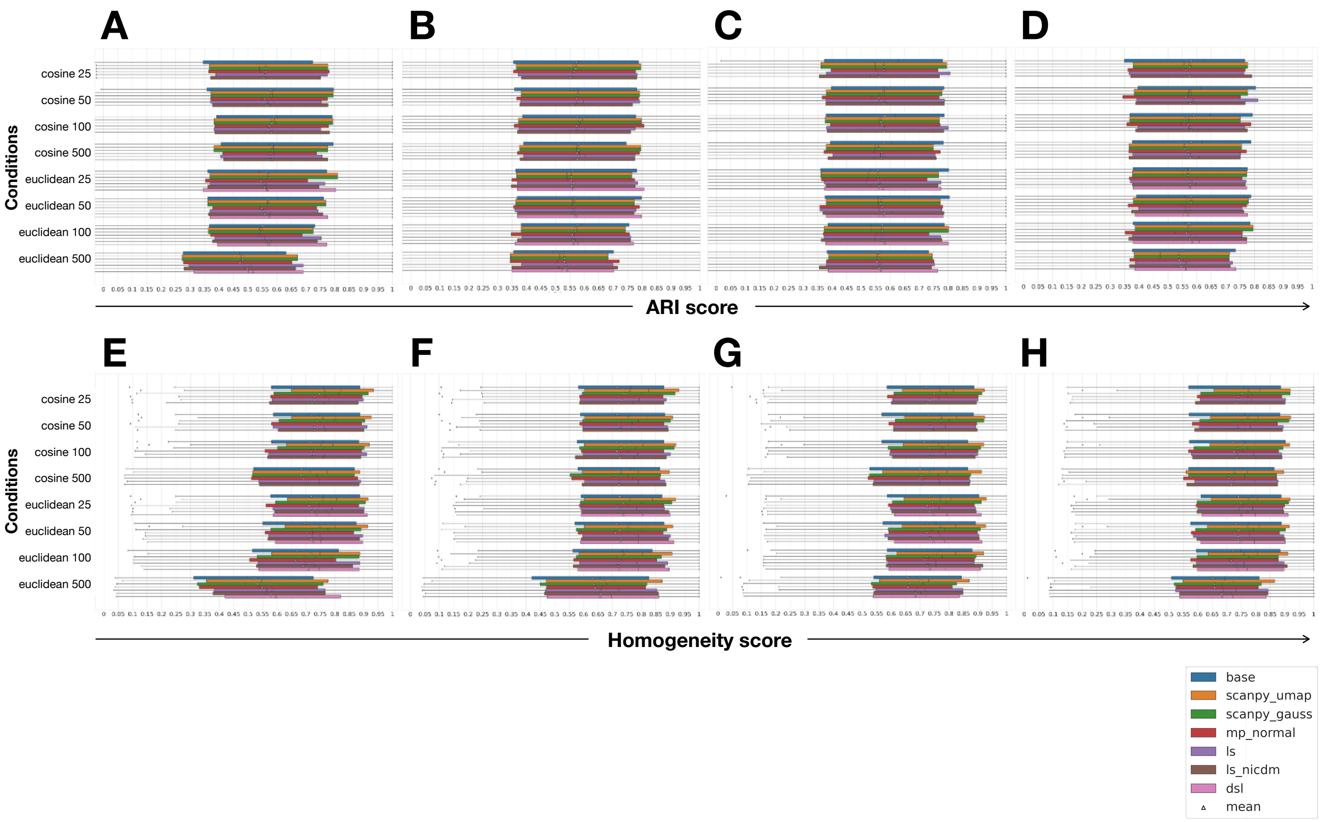


Supplementary Figure 10: clustering scores done with the Seurat **(A,B,E,F)** or Duo **(C,D,G,H)** preprocessing with **(A,C,E,G)** or without **(B,D,F,H)** scaling, for low-ID datasets: ARI scores **(A,B,C,D)** and homogeneity scores **(E,F,G,H)**.

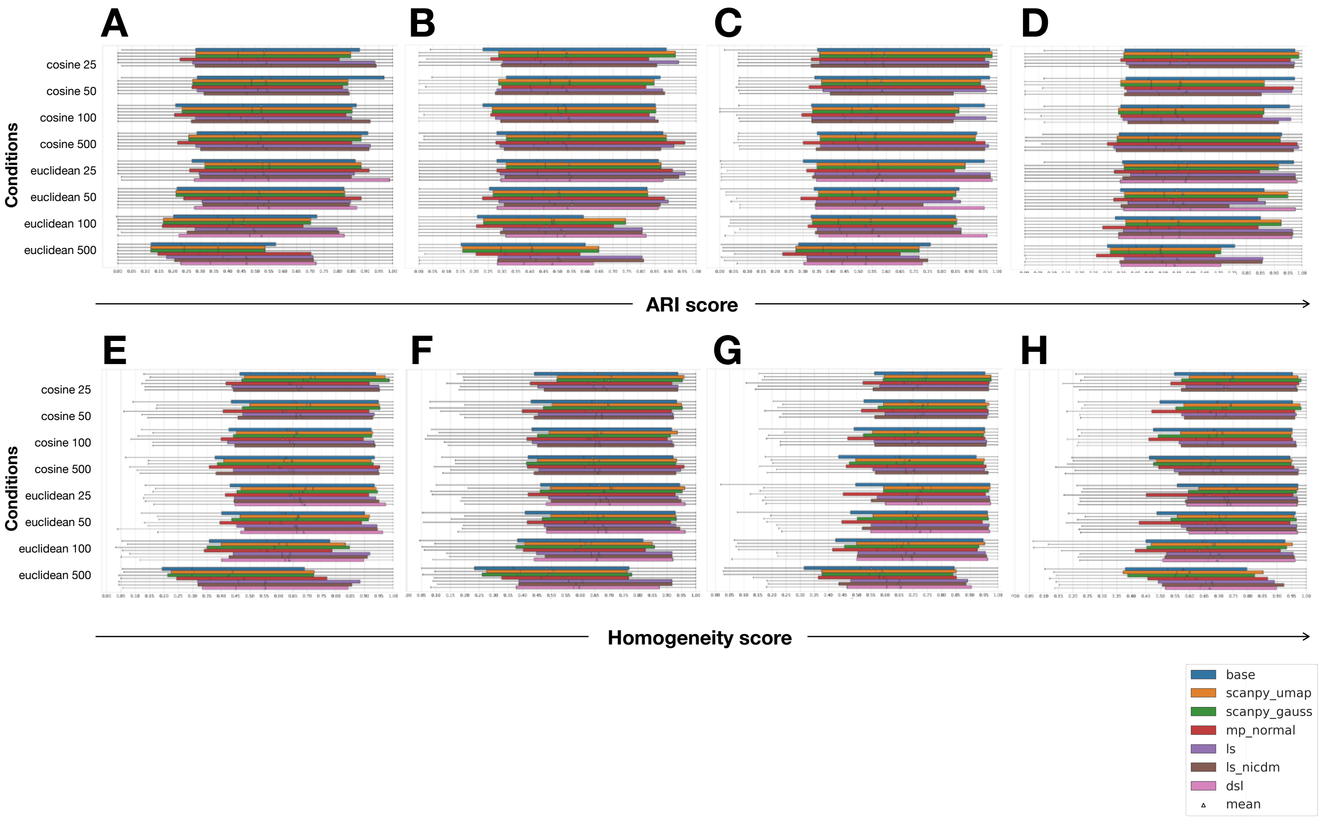


Supplementary Figure 11: clustering scores done with the Seurat **(A,B,E,F)** or Duo **(C,D,G,H)** preprocessing with **(A,C,E,G)** or without **(B,D,F,H)** scaling, for gold-standards datasets: ARI scores **(A,B,C,D)** and homogeneity scores **(E,F,G,H)**.

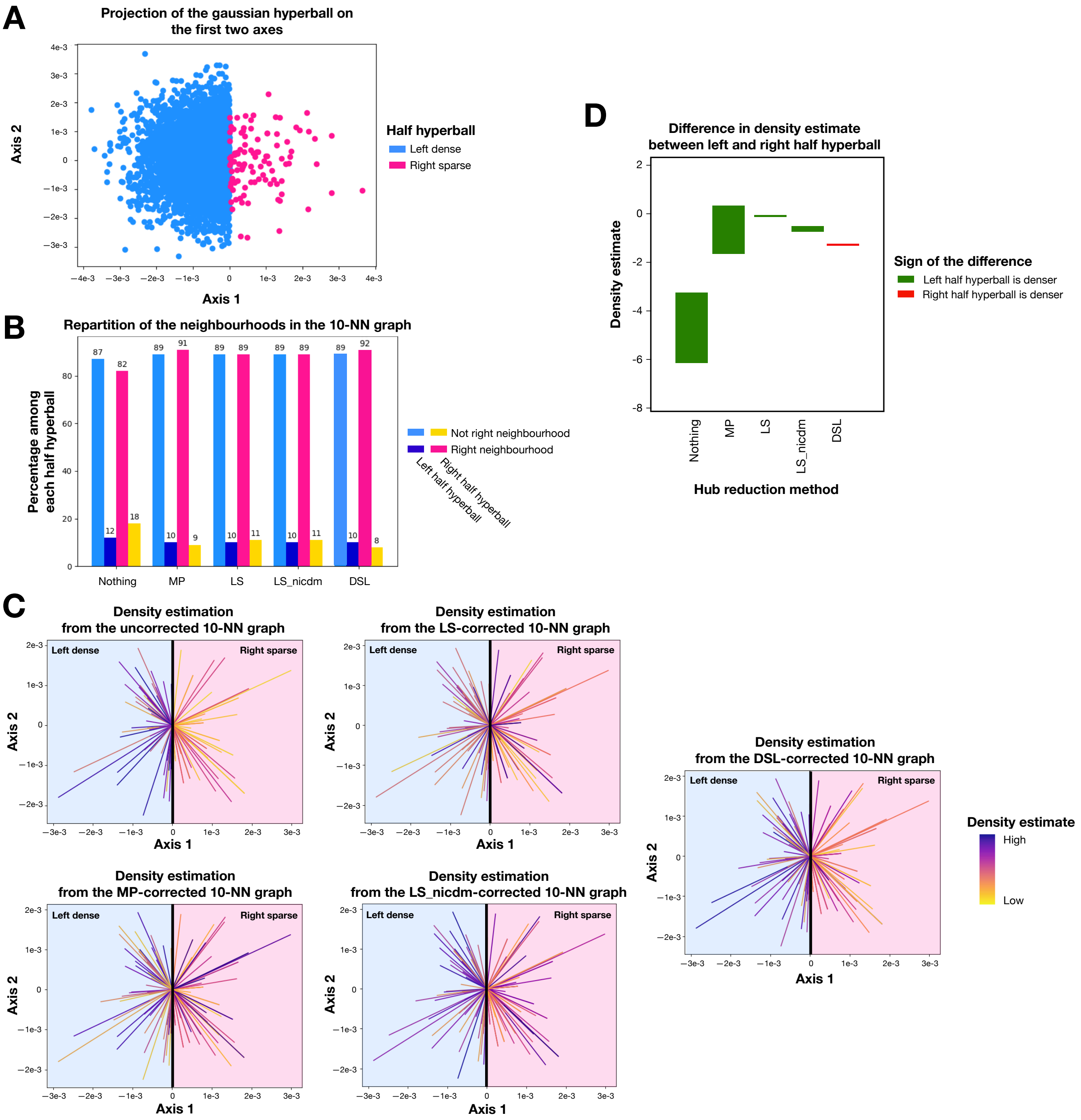


Supplementary Figure 12: Correcting heterogeneous densities with hubness reduction. **(A)** Gaussian ball in 10 dimensions, with 5000 points in the half hyperball left of the hyperplan x=0 and 100 points in the right half hyperball, projected on the first two dimensions. **(B)** Proportion of points of each half hyperball in the neighborhood of the right half hyperball. **(C)** Visualization of the density estimate calculated from the unweighted k-NN graph before and after hubness reduction. The source of the density estimate is at the center of the Gaussian ball and the targets are picked randomly in each half hyperball. Each line connect the source and a target and its color represents the density estimate. The pale background colors represent the two half spaces: blue for the left one, pink for the right one. **(D)** Difference in the density estimates between the left and right half hyperballs. Each edge of a bar is the mean density estimate in one of the half hyperballs. If the rectangle is green, the lower border is the estimate from the right half hyperball; if it is red, it is from the left one.

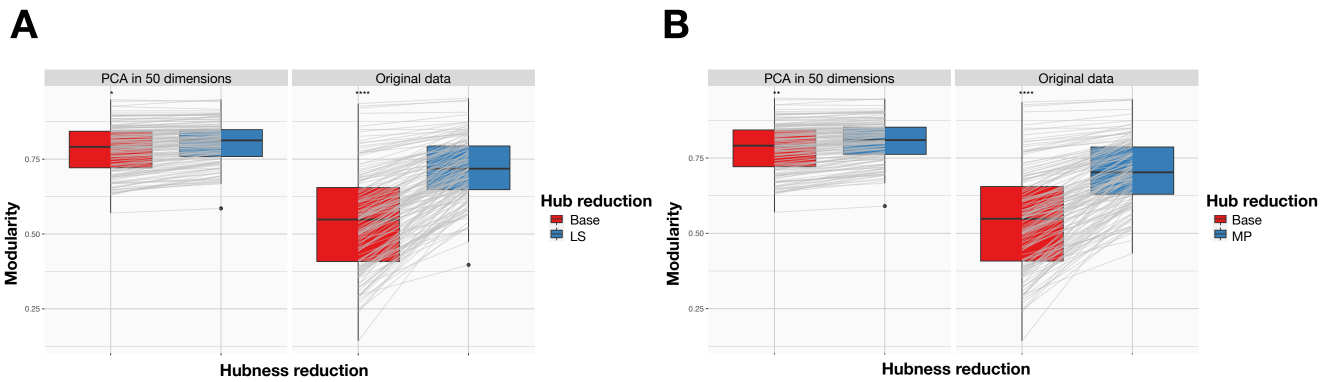


Supplementary Figure 13: Modularity improvement upon hubness reduction. Per-dataset modularity of the Louvain clustering with (left panel) or without PCA (right panel). Comparison between the modularity with or without hubness reduction, performed with the LS **(A)** or the MP algorithm **(B)**.

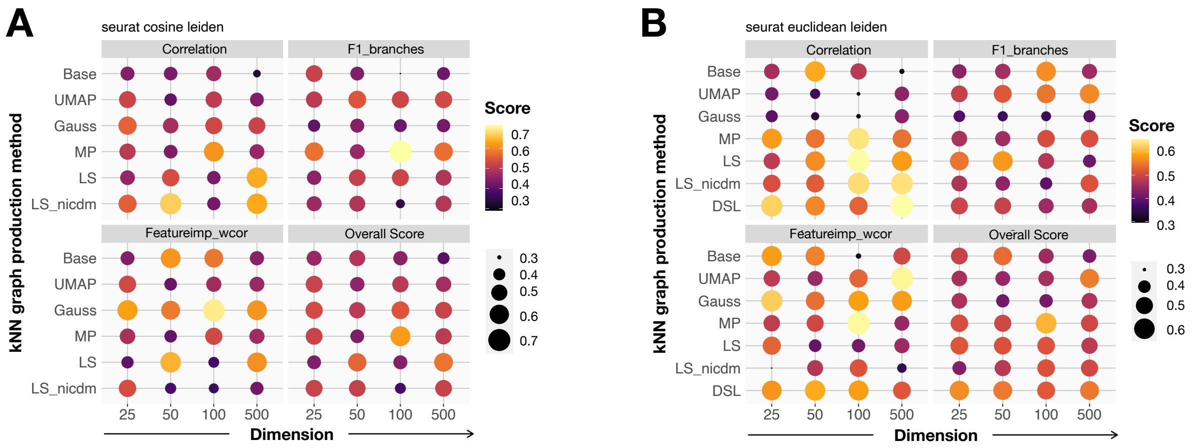


Supplementary Figure 14: Averaged trajectory inference scores on low-ID datasets, using the Seurat recipe. **(A)** Mean TI quality metrics with the cosine dissimilarity and the Leiden clustering. **(B)** Mean TI quality metrics with the Euclidean metric and the Leiden clustering.

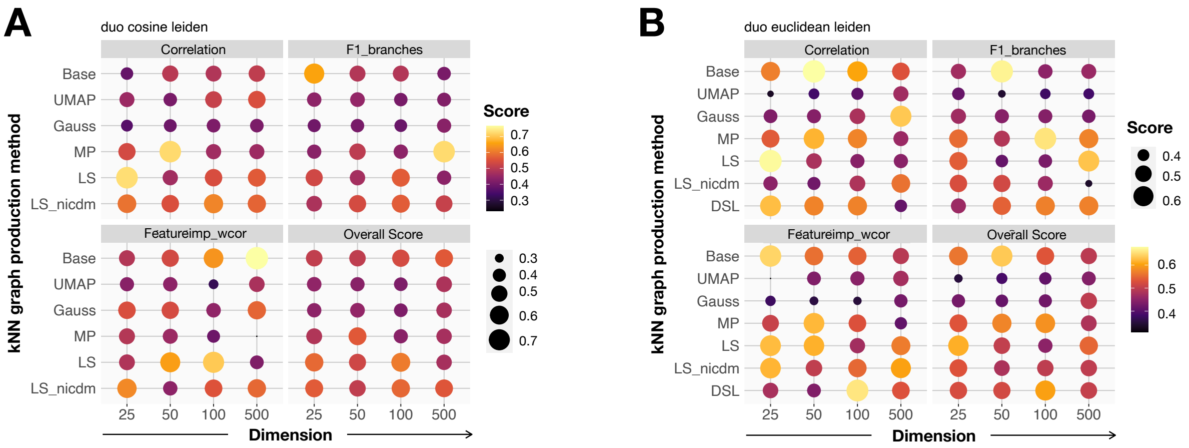


Supplementary Figure 15: Averaged trajectory inference scores on low-ID datasets, using the Duo recipe. **(A)** Mean TI quality metrics with the cosine dissimilarity and the Leiden clustering. **(B)** Mean TI quality metrics with the Euclidean metric and the Leiden clustering.

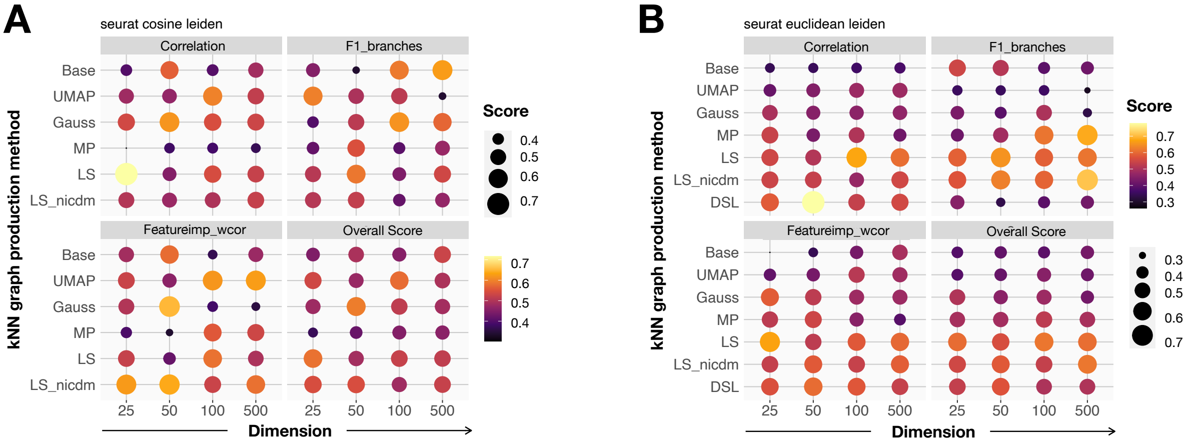


Supplementary Figure 16: Averaged trajectory inference scores on high-ID datasets, using the Seurat recipe. **(A)** Mean TI quality metrics with the cosine dissimilarity and the Leiden clustering. **(B)** Mean TI quality metrics with the Euclidean metric and the Leiden clustering.

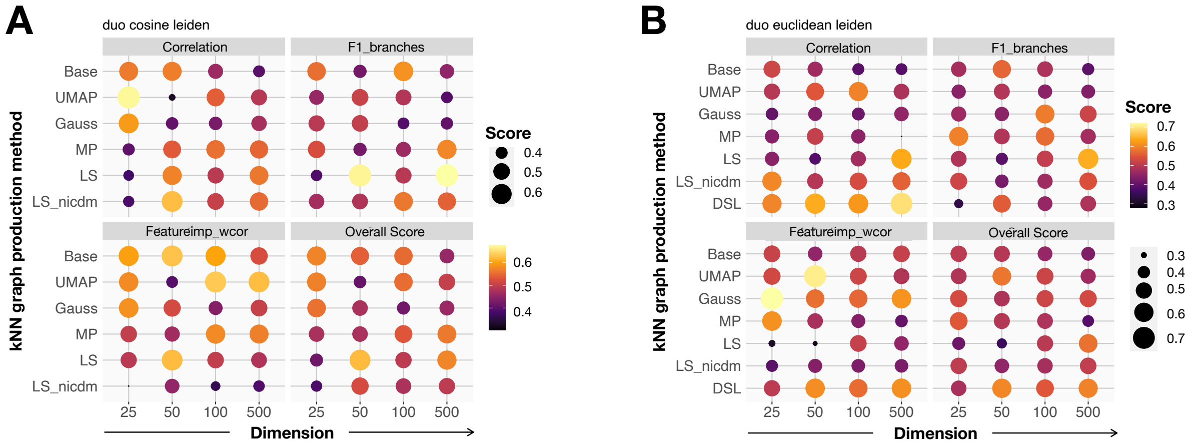


Supplementary Figure 17: Averaged trajectory inference scores on high-ID datasets, using the Duo recipe. **(A)** Mean TI quality metrics with the cosine dissimilarity and the Leiden clustering. **(B)** Mean TI quality metrics with the Euclidean metric and the Leiden clustering.

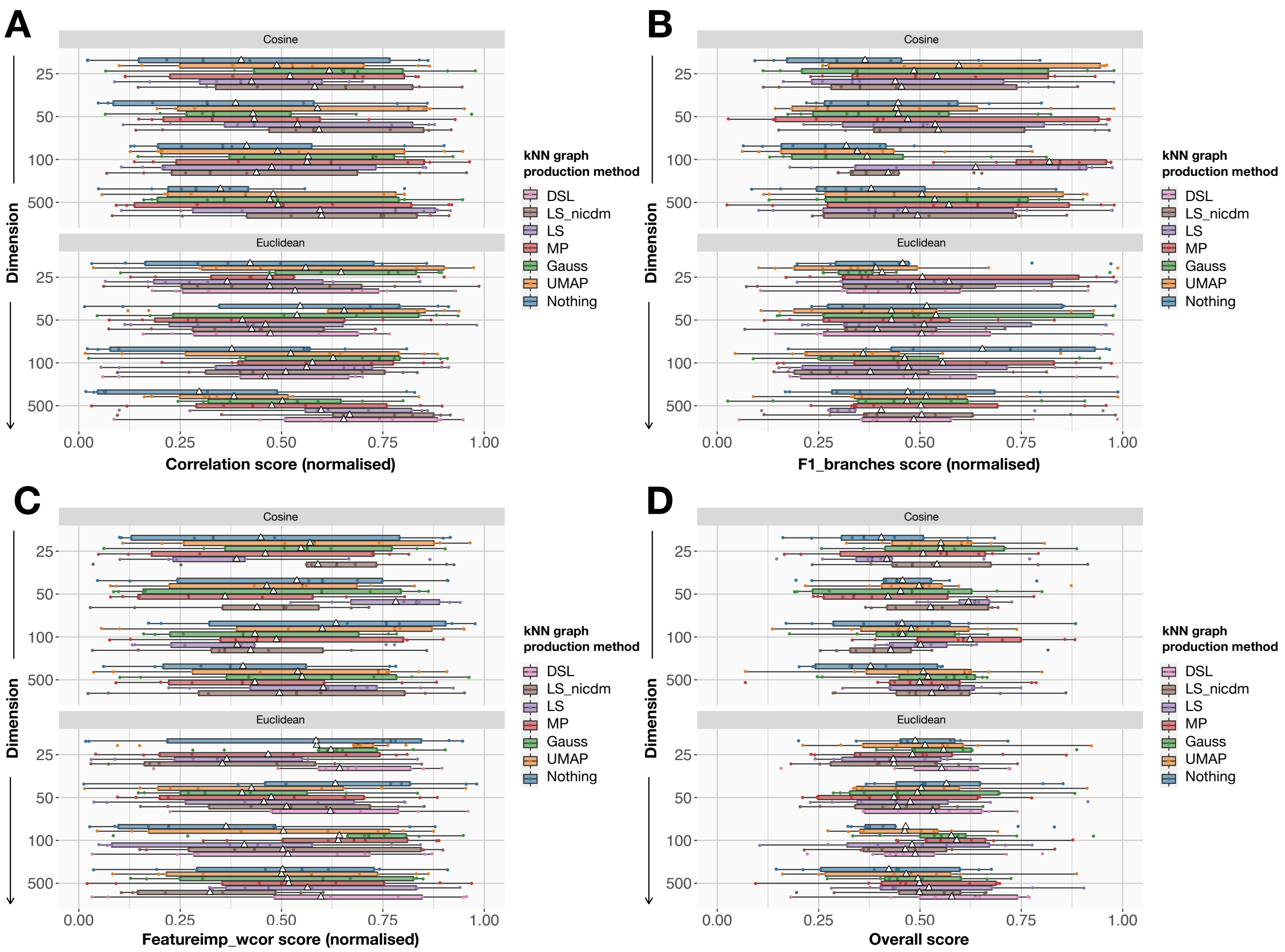


Supplementary Figure 18: Per-dataset trajectory inference scores on low-ID datasets, using the Seurat recipe and Leiden clustering. **(A)** Detailed correlation scores. **(B)** Detailed F1_branches scores. **(C)** Detailed featureimp_wcor scores. **(D)** Detailed overall score.

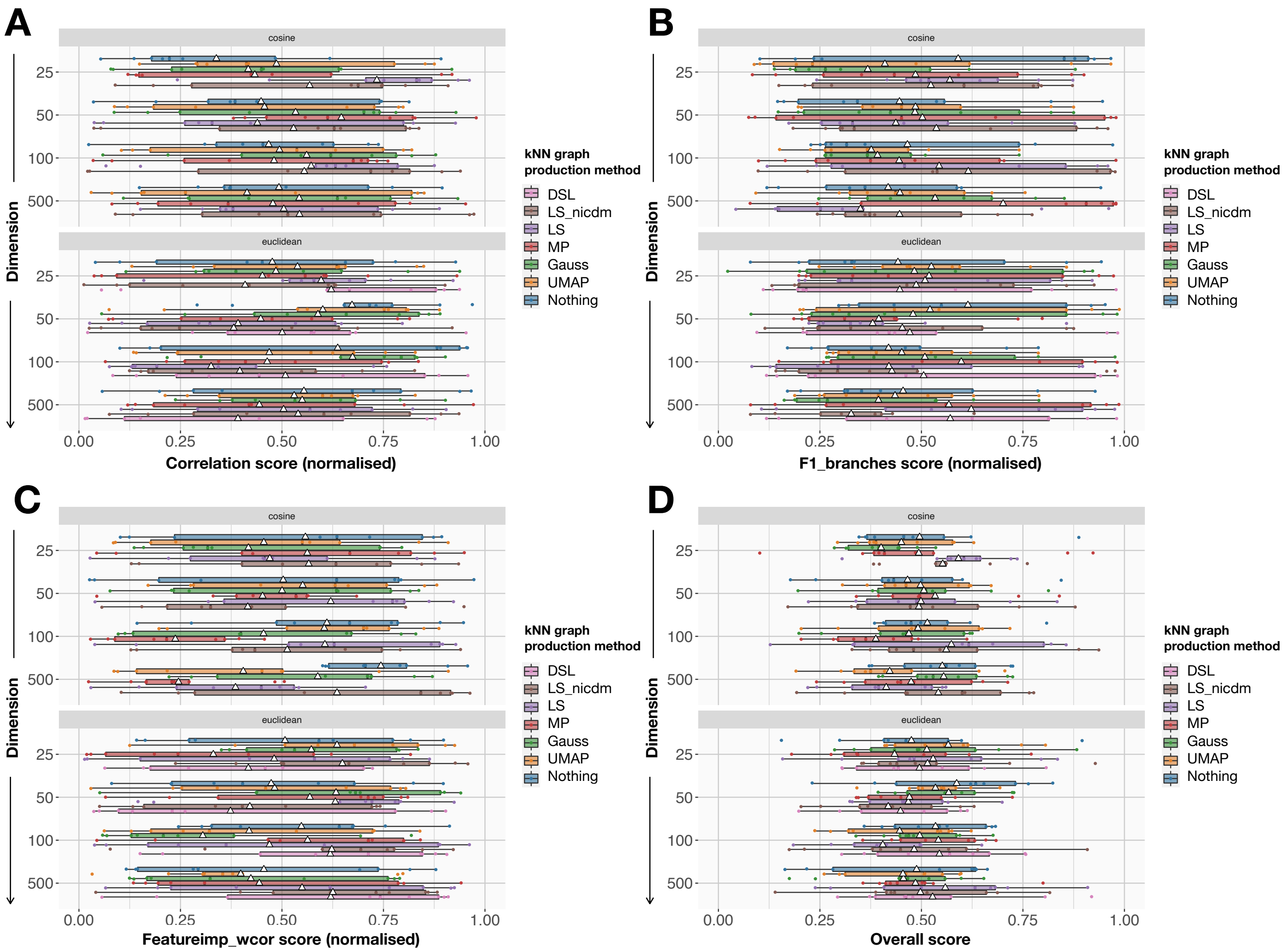


Supplementary Figure 19: Per-dataset trajectory inference scores on low-ID datasets, using the Duo recipe and Leiden clustering. **(A)** Detailed correlation scores. **(B)** Detailed F1_branches scores. **(C)** Detailed featureimp_wcor scores. **(D)** Detailed overall score.

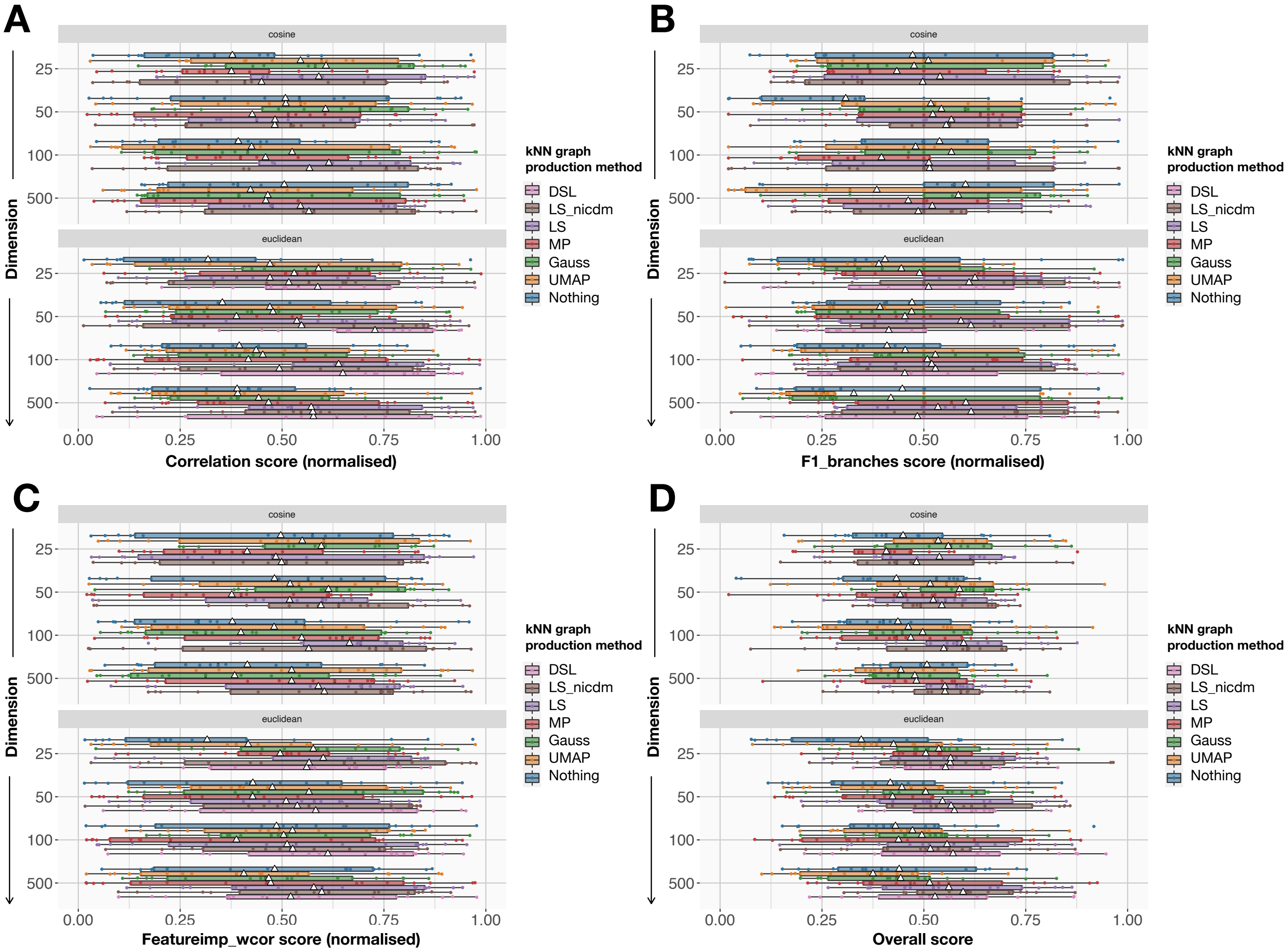


Supplementary Figure 20: Per-dataset trajectory inference scores on high-ID datasets, using the Seurat recipe and Leiden clustering. **(A)** Detailed correlation scores. **(B)** Detailed F1_branches scores. **(C)** Detailed featureimp_wcor scores. **(D)** Detailed overall score.

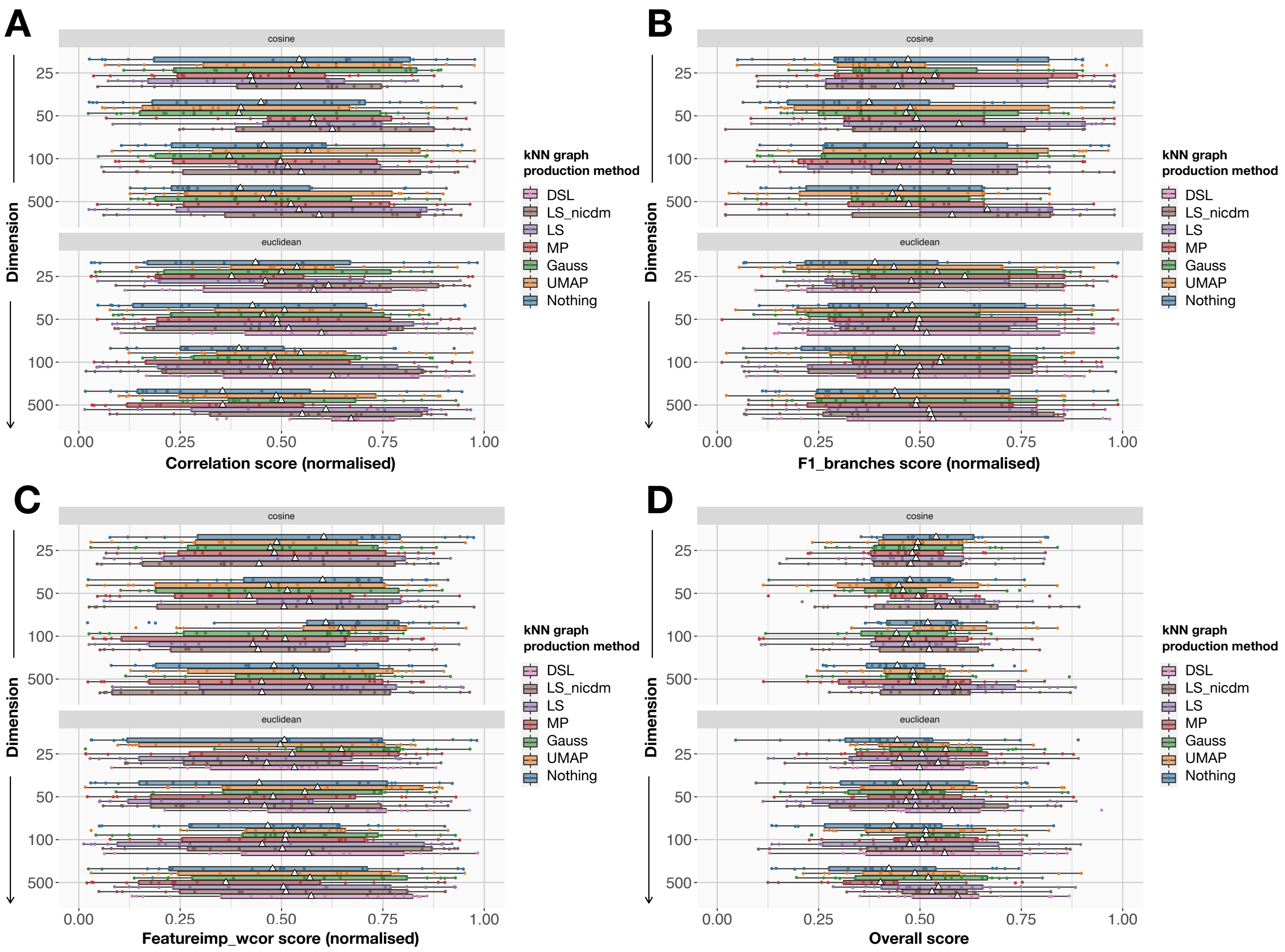


Supplementary Figure 21: Per-dataset trajectory inference scores on high-ID datasets, using the Duo recipe and Leiden clustering. **(A)** Detailed correlation scores. **(B)** Detailed F1_branches scores. **(C)** Detailed featureimp_wcor scores. **(D)** Detailed overall score.

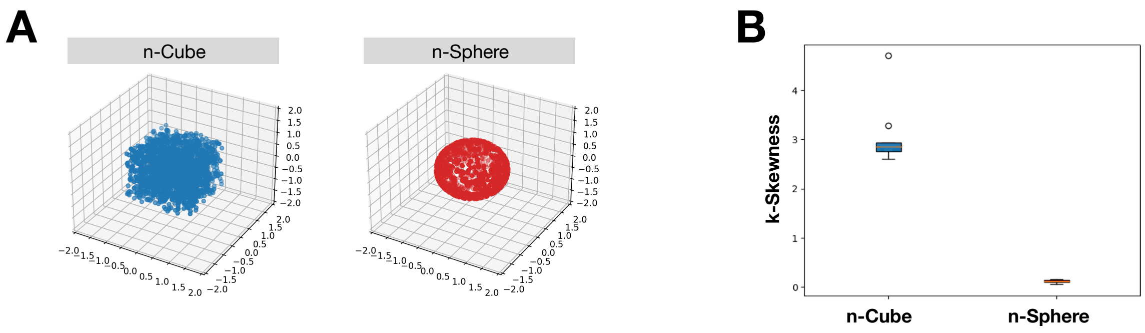


Supplementary Figure 22: n-Cube and n-Sphere. **(A)** 3D plot of a 3-Cube (left) and a 3-Sphere (right). **(B)** 5k-Skewness of a 50-Cube and a 50-Sphere, each containing 5,000 points, and k=50, computed 10 times.

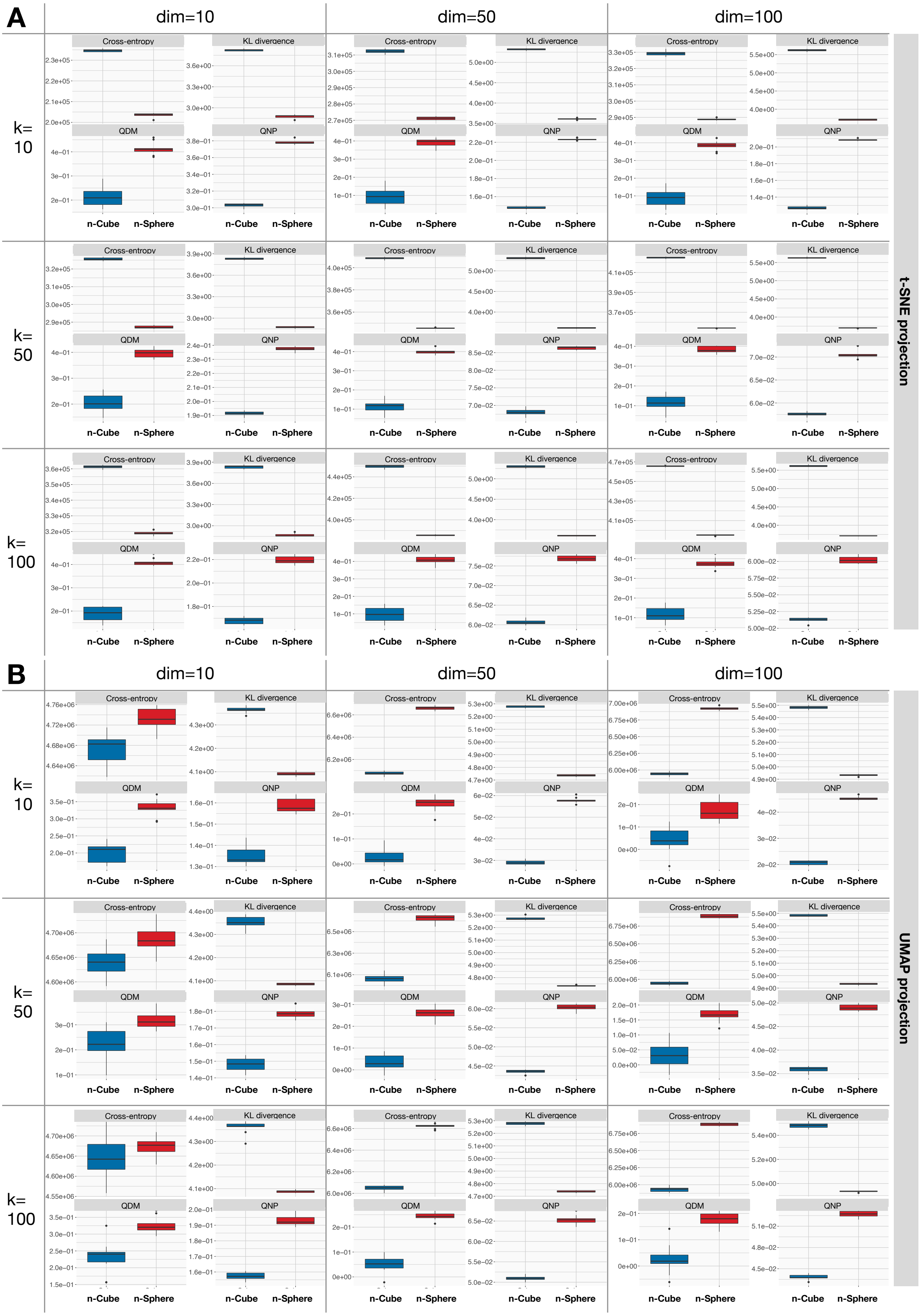


Supplementary Figure 23: Visualisation quality metrics of the n-Cube and the n-Sphere. We show the $n$-cube and the $n$-sphere after t-SNE **(A)** or UMAP **(B)** projections, with different values for the size of the neighborhood $k$ and the number of dimensions $n$

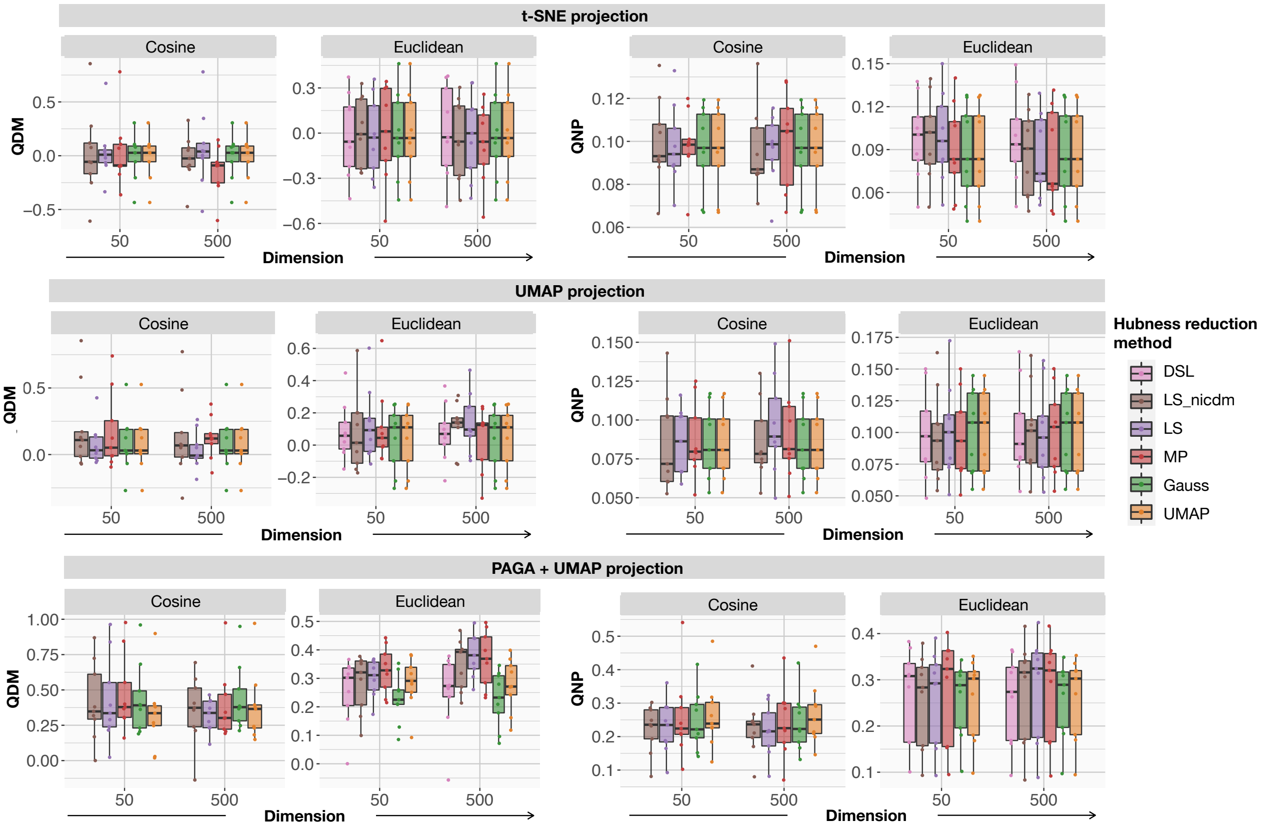


Supplementary Figure 24: QDM and QNP before or after hubness reduction, evaluated after various visualisation algorithms, compared to the PCA with 500 PCs, for low-ID datasets. We project low-ID datasets either with t-SNE (top row), UMAP (middle row) or PAGA+UMAP (bottom row) and evaluate QDM (left column) and QNP (right column). The different projections are computed either with the cosine dissimilarity or the Euclidean metric, and using the two Scanpy k-NN graphs or the four hub-reduced graphs.

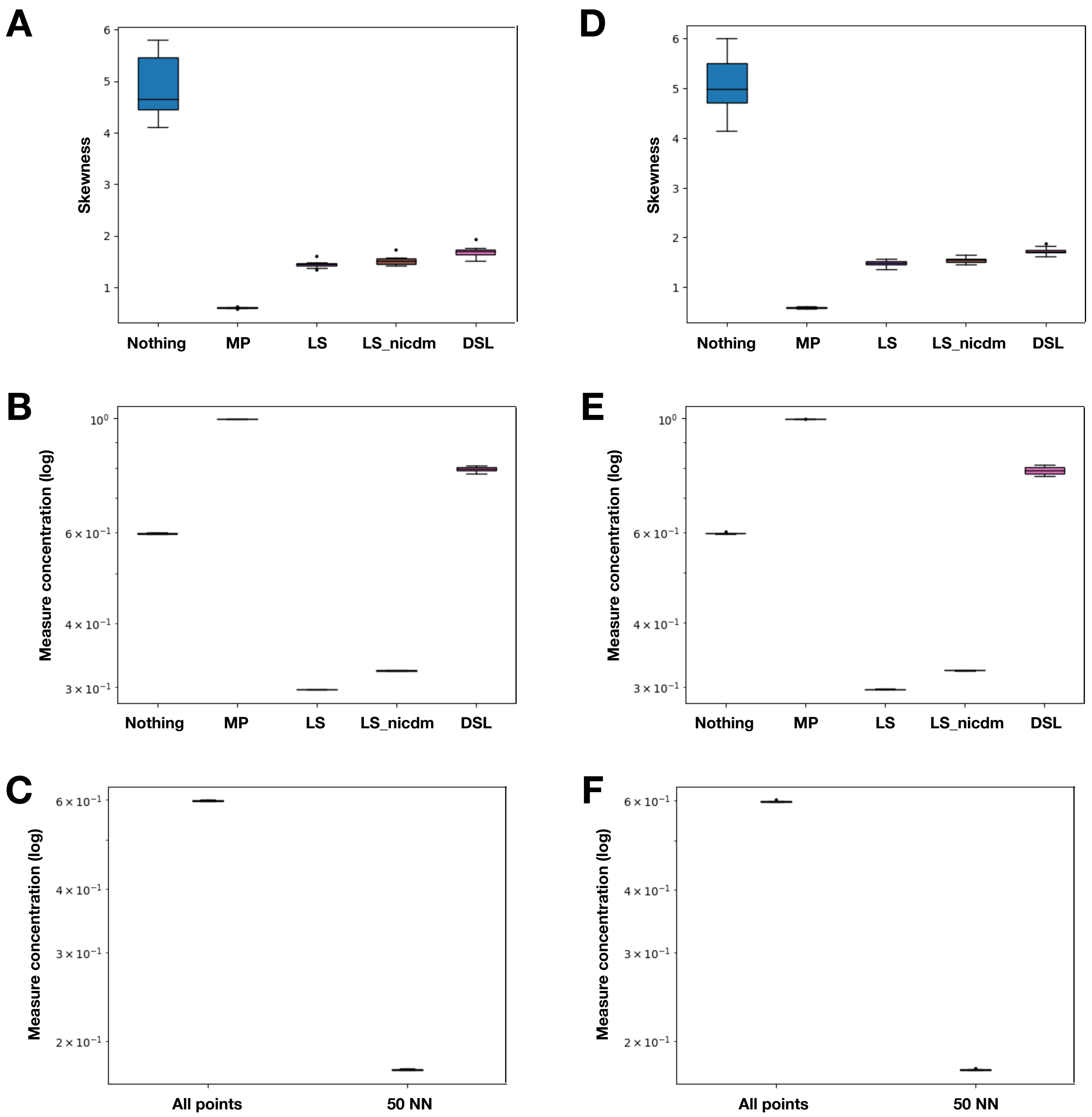


Supplementary Figure 25: Evaluation of the impact of hubness correction on hubness and measure concentration. We computed Gaussian distributions, either a single blob **(A,B,C)** or two distinct blobs **(D,E,F)**. Skewness of the data with or without hubness reduction **(A,D)**. Global measure concentration with or without hubness reduction **(B,E)**. Measure concentration without hubness reduction, either considering all points, or the 50 nearest neighbors **(C,F)**
